# Integration of Mouse and Human Single-cell RNA Sequencing Infers Spatial Cell-type Composition in Human Brains

**DOI:** 10.1101/527499

**Authors:** Travis S Johnson, Zachary B Abrams, Bryan R Helm, Peter Neidecker, Raghu Machiraju, Yan Zhang, Kun Huang, Jie Zhang

## Abstract

Technical advances have enabled the identification of high-resolution cell types within tissues based on single-cell transcriptomics. However, such analyses are restricted in human brain tissue due to the limited number of brain donors. In this study, we integrate mouse and human data to predict cell-type proportions in human brain tissue, and spatially map the resulting cellular composition. By applying feature selection and linear modeling, combinations of human and mouse brain single-cell transcriptomics profiles can be integrated to “fill in” missing information. These combined “in silico chimeric” datasets are used to model the composition of nine cell types in 3,702 human brain samples in six Allen Human Brain Atlas (AHBA) donors. Cell types were spatially consistent regardless of the scRNA-Seq dataset (91% significantly correlated) or AHBA donor (p-value = 4.43 ×10^-20^ by t-test) used in the model. Importantly, neuron nuclei location and neuron mRNA location were correlated only after accounting for neural connectivity (p-value = 1.26×10^-10^), which supports the notion that gene expression is a better indicator than nuclei location of cellular localization for cells with large and irregularly shaped cell bodies, such as neurons. These results advocate for the integration of mouse and human data in models of brain tissue heterogeneity.

## Introduction

An important goal for neuroscience is to understand how structural, anatomic, and cellular heterogeneity relate to brain development, cognitive function, neurological disease, and senescence/degeneration [1]. Recently, over 40 cell types with unique anatomic characteristics [2] as well as complex spatially distinct anatomic connections [3] were identified in the mouse brain. Due to the spatial and functional complexity of these brain structures, there is a strong need to combine anatomically explicit tissue sampling with functional assessment to “localize molecularly defined subtypes in tissues, with simultaneous detection of morphology, activity, or connectivity” [4]. Fortunately, a confluence of enhanced gene expression sequencing capabilities, deposition of well-curated sample data into publicly available repositories, and improved analytical tools can help address these gaps in our existing knowledge of cellular and functional heterogeneity in the brain.

Single-cell sequencing has empowered a new generation of transcriptomics-based cell classification studies,[2, 5, 6]. Single-cell-resolution transcriptomes have enabled the classification of cell type and function based on similarities and dissimilarities in gene expression, which in turn has identified novel and functionally distinct subtypes of neuronal and glial cells [2, 7–9]. These findings suggest that brain cells are more functionally diverse than previously assumed. Moreover, these cell types can be predicted and identified based on gene expression and evaluated with existing knowledge about their role within the brain. To date, this approach has not been utilized to estimate the tissue heterogeneity of spatially sampled and anatomically distinct human brain tissues, though such analyses have been conducted for other human organs and tissues [10–12].

The Allen Human Brain Atlas (AHBA) is a database that curates transcriptomics data from human brain tissues while also providing additional sample information, including spatial and anatomic locations [13]. One major barrier to studying anatomically explicit gene expression patterns in human brains is the paucity of relevant donor samples [14]. Previous studies have integrated disparate datasets with the AHBA to supplement the low numbers of human brain donors and harness features unique to each dataset [15]. The AHBA database therefore provide a unique opportunity to model cellular heterogeneity spatially across multiple brains and anatomic regions. One limitation, however, is that samples can be highly variable and biased by the donor’s age, cause of death, and time postmortem. For this reason, it is important to leverage additional information, where possible, to complement the existing data infrastructure for the purposes of large-scale analyses.

One potential solution for augmenting information from the AHBA is to incorporate data from the brains of other well-studied model organisms. It has already been shown that mouse and human interneurons share similar co-expression motifs [16] and that brain region-specific gene expression is homologous between human and mouse [18]. In addition, feature selection can correct for some interspecies variance [17].

In this manuscript, therefore, we address several key challenges that are not only important for neurological research but also for medical computational modeling, including the integration of mouse and human data sets, prediction of brain cell-type composition from expression data, and accurate mapping of anatomical location and spatial heterogeneity. First, we propose methods for integrating scRNA-Seq data from mouse and human brain tissues to estimate cell-type composition in heterogeneous brain tissue. Second, we validate our methods on simulated samples generated from scRNA-Seq data. Third, we estimate the cell-type composition in spatially sampled human brains. Finally, we show that neural connectivity (e.g., neuron size, axonal projection size) alters the RNA composition in brain tissue. Our aims were as follows:

**Aim 1:** Create an analysis pipeline that accurately and precisely integrates human and mouse data to better predict cell-type composition based on gene expression.

**Aim 2:** Use integrated data to predict cell-type composition in unknown human samples that are spatially distributed (i.e., anatomically explicit) throughout the human brain.

**Aim 3:** Examine whether cell types correspond with known brain data to the 1) anatomical location in the brain; 2) neural connectivity; and 3) nuclear density of the cells.

Previously, to obtain cell-type-specific RNA profiles, researchers physically sorted cells before sequencing or computationally sorted samples after sequencing using expression deconvolution (**Figure 1A-1** [19], A-2 [20]). Deconvolution is the process by which homogenous elements are estimated from a heterogeneous mixture. This new paradigm (Figure 1A-3) is complementary to the above two workflows, and is adopted in this study. Specifically, single cells are sequenced individually then stratified computationally after clustering into groups, which are then used as input for deconvolution. The general mathematical estimation of homogeneous tissue proportions in heterogeneous tissue mixtures has already been described in detail [19, 21–29].

**Figure 1.**
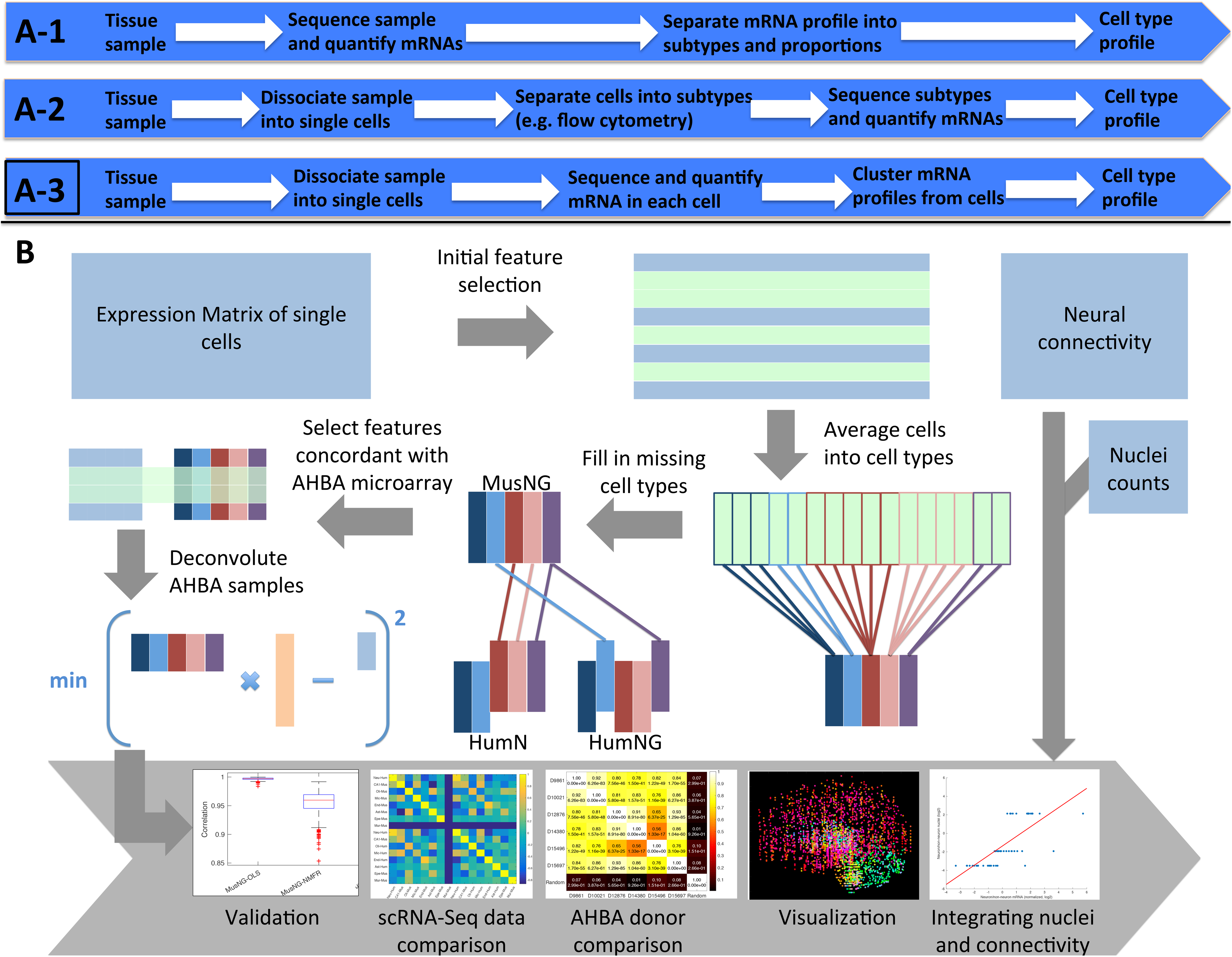
The classic characterization of cell types from RNA-Seq data. A-1. and A-2. were used before the advent of scRNA-Seq. A-3. is the current state of the art, and was chosen as the basis for characterization in this study. B. The workflow of this study from raw scRNA-Seq datasets (light blue) to processed data and downstream analyses.

Although deconvolution methods using both cell-type proportions and cell-type expression profiles have been studied previously using microarray data [19, 21, 23–30], there is limited research on the deconvolution method applied to microarray or bulk RNA-Seq samples using profiles derived from scRNA-Seq [10, 31]. Furthermore, the combination of mouse and human datasets in this process is limited to species-specific deconvolution of pancreatic tissue to identify biomarkers [10]. The method presented in this paper expands these general ideas to the brain, and combines human and mouse expression profiles prior to deconvolution.

To address our first aim, we integrated mouse brain cell expression information to complete various sets of human brain cell types. To address our second aim, we inferred cell-type composition from “in silico chimeric” mouse-human data sets and mapped these characterizations anatomically in human brain samples. To address aim 3, cell types were evaluated against their anatomic localization pattern, and neuron nuclei density was compared with neuron composition across anatomic locations. This workflow required surmounting several challenges, including:

1. Implementation of a robust deconvolution method and validation (Aim 1);
2. Integration of mouse and human expression data (Aim 1);
3. Integration of scRNA-Seq and microarray data for deconvolution in a broad set of scenarios (Aim 2);
4. Integration of RNA-Seq with the spatial mapping of cell type and neural connectivity (Aim 3).

Importantly, due to the large size and irregular shape of neurons and other neural cells, the relationships between RNA, nuclei, and cell shape are critically important for modeling cells from brain tissue. We believe that the nuclei density or mRNA quantity alone may not be an optimal representation of cellular localization in large and irregularly shaped cell types like neurons. The Allen Brain Institute provides rich information about brain connectivity from mouse brain tissues by labeling axonal projections with rAAV tracers and imaging with two-photon tomography. The results are neural projection volumes at both the injection and target sites. By integrating mouse brain connectivity data [3], we show that these projections may affect cell-type RNA quantities across the brain spatially, resulting in changes to cell-type quantification. By incorporating neural connectivity information into the deconvolution process, we show our deconvolution results replicate neural nuclei location.

## Results

### Cell-type proportion estimates are accurate for simulated scRNA-Seq tissue

The workflow of the experimental process is shown in Figure 1. The cell-type proportions (i.e., composition) that resulted from deconvolution of each of the three scRNA-Seq datasets were highly correlated with the underlying true proportions of single cells (**Figure 2**). We observed that the true proportions could be reproduced from the trained model for any of the original scRNA-Seq datasets (Figure 2A). Furthermore, the ordinary least squares (OLS) model always performed better than the non-negative matrix factorization regression (NMFR) model based on the Pearson’s correlation coefficient (PCC); the OLS model was therefore used for the remainder of the study (Figure 2B). In general, the deconvolution parameters and the gene variation among the datasets and samples do not affect the accuracy of the model. The accuracy decreased the most for a step size of less than ~0.2 (Supplementary Figure S3); we therefore used 0.3 as the step size and 0.9 as the standard deviation constant based on a grid search (Supplementary Figure S3).

**Figure 2.**
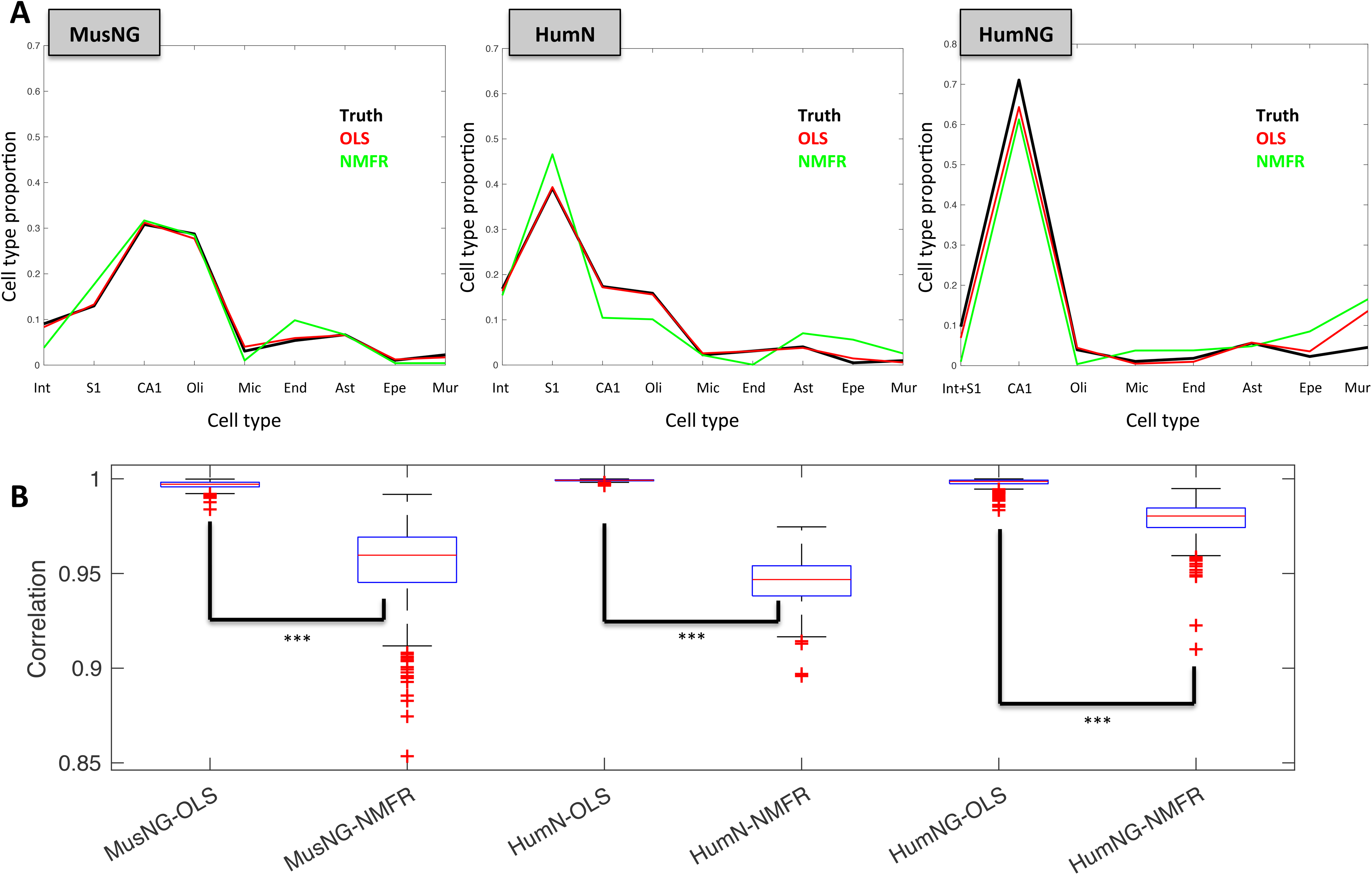
A comparison of cell-type estimate accuracy using the data from each of the three scRNA-Seq datasets. **A.** Representative examples of the cell-type proportions using each scRNA-Seq dataset on donor 10021. MusNG: A MusNG simulated sample deconvoluted using the cell types from MusNG. HumN: A HumN simulated sample deconvoluted using the cell types from HumN. HumNG: A HumNG simulated sample deconvoluted using the cell types from HumNG. **B.** Pearson’s correlation coefficients (PCC) between the true proportions of cell types from simulated tissue samples and predicted proportions. The experiments were repeated 100 times for each scRNA-Seq dataset, regression method, and Allen Human Brain Atlas (AHBA) donor. ***indicates p-value < 0.001.

### Cell-type proportion estimates of AHBA donors are consistent across scRNA-Seq datasets

The cell-type proportions were estimated in each of the AHBA donors using scRNA-Seq data (i.e., MusNG, HumN, and HumNG; see Supplementary Figure 1 and Materials and Methods for details). The resulting estimated cell-type proportions (three per AHBA donor) were consistent regardless of the scRNA-Seq dataset, and were used to deconvolute data from each donor (Figure 3A).

**Figure 3.**
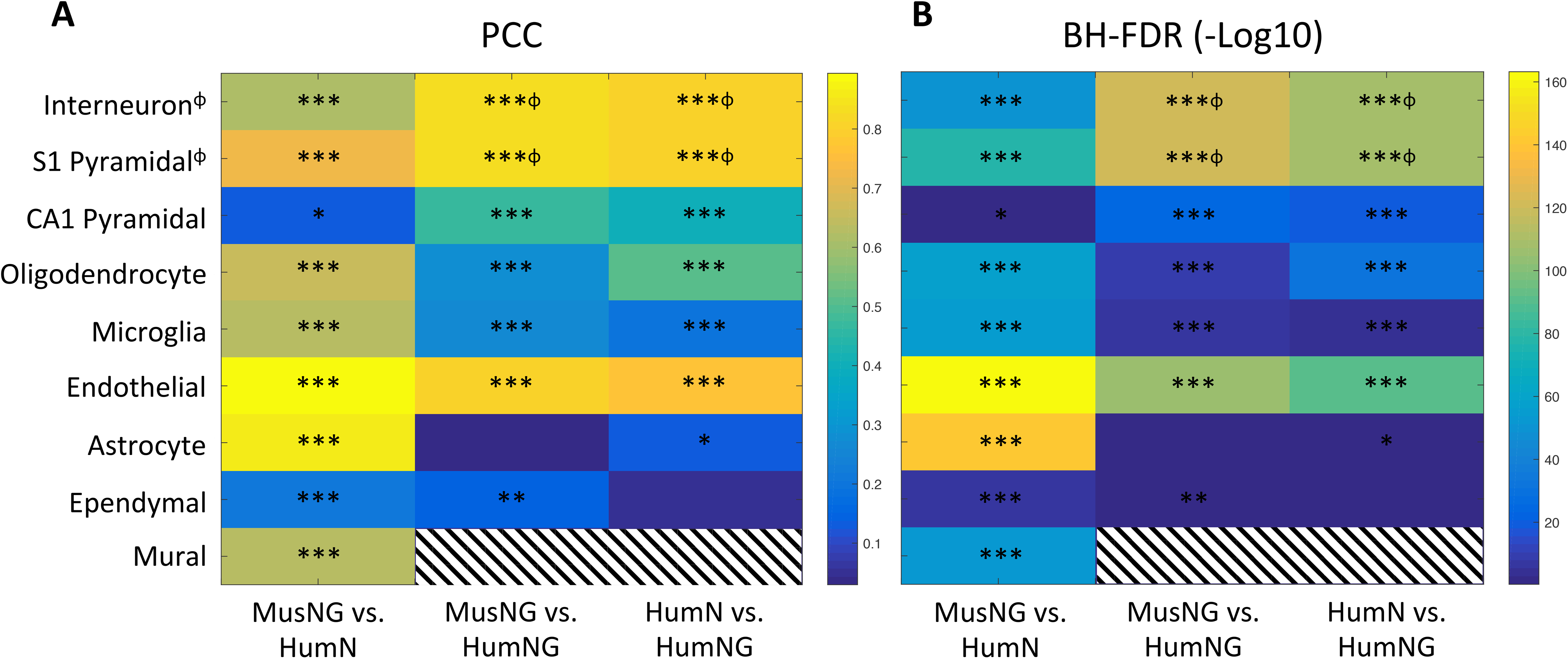
Pearson’s correlation coefficients (PCC) and Benjamini-Hochberg false discovery rates (BH-FDR) for each cell-type prediction between each pair of the three input datasets. **A.** PCC values for each cell-type correlation across all samples deconvoluted with different scRNA-Seq datasets. **B.** Negative log10 BH-FDR values for the PCC correlations in A. The scale is the exponent of the BH-FDR. *indicates correlations p < 0.05, **p < 0.01, *** p < 0.001. ϕ indicates the two cell types that were combined for the MusNG+HumNG and HumN+HumNG comparison such that the first two rows in the right two columns in A and B are the same value. See Supplemental Figure 1 for more information. Stripes indicate that there were not enough data to generate a correlation.

We expect cell-type correlations among datasets to be positive, which would suggest that the cell-type proportions are consistent between input datasets (Figure 3A). We observed that 91% (21/23) of the correlations were significantly positively correlated (Figure 3A) with a low Benjamini-Hochberg false discovery rate (BH-FDR) (Figure 3B). Neural cell-type correlations were much stronger than glial cell-type correlations because glial cell types were harder to differentiate (Supplementary Figure S4-10). Furthermore, we found that the same cell-type correlations were positively correlated more frequently than mismatched cell-type correlations (Supplementary Figure S4-10, Supplementary Table II). The cell-type estimates using the MusNG dataset were most similar to the HumN datasets for AHBA donors. The HumN versus HumNG correlations were higher than the MusNG versus HumNG correlations despite the same feature set in the HumN and MusNG datasets (Figure 3, Supplementary Figure S4-10). This finding is caused by the gene expression profile similarity between the HumN and HumNG datasets. Within each of these comparisons, there were also indications that dissimilar cell types follow similar patterns to other cell types between datasets. For example, human oligodendrocyte proportions correlate with mouse astrocyte proportions (Supplementary Figure S4-10B,D,F). This finding could be attributed to similar glial expression profiles or an overlap between the two cell types in anatomical space. We observed that the absolute proportions of neural cell types, especially pyramidal cells, were much higher than glial or interneuron cell types (Supplementary Figure S4-10A,C,E) in cerebrum samples. This finding may be attributed to the higher quantity of neural cell mRNA. Alternatively, neural cell mRNA may have a specific signature that is easier to detect.

### Cell-type proportion spatial mappings are consistent among AHBA donors

We demonstrated that scRNA-Seq datasets were not sources of bias by comparing the cell-type proportions using different scRNA-Seq datasets. This lack of influence of the scRNA-Seq dataset indicates that our feature selection and deconvolution method is effective. Therefore, we next show that the donor is not a major contributor of variance to the cell-type proportions.

The estimated cell-type proportions for each donor were mapped to the anatomical location from which they were sampled. For each anatomic location, the mean cell-type proportion was used to produce an aggregated cell-type proportion for each anatomic location. By using scRNA-Seq data to calculate the cell-type proportions for each anatomic location, we observe high cell-type consistency between donors for each of the anatomic locations: the average pairwise PCC was 0.87 across all three scRNA-Seq datasets (**Figure 4**). The PCC among AHBA donors was also significantly higher and significantly more positively correlated than randomly shuffled data (Figure 4, p-values MusNG: 1.73 ×10^-23^, HumN: 1.92 ×10^-27^, HumNG: 1.18 ×10^-14^ by t-test; averaged: 4.43 ×10^-20^). Furthermore, none of the randomized comparisons were significantly correlated to every non-random comparison (Figure 4, Supplementary Figure S11-13). We also observed variability among brain donors; for example, AHBA donor 9861 had significantly lower correlations than other donors (p-value = 1.46 ×10^-7^ by t-test).

**Figure 4.**
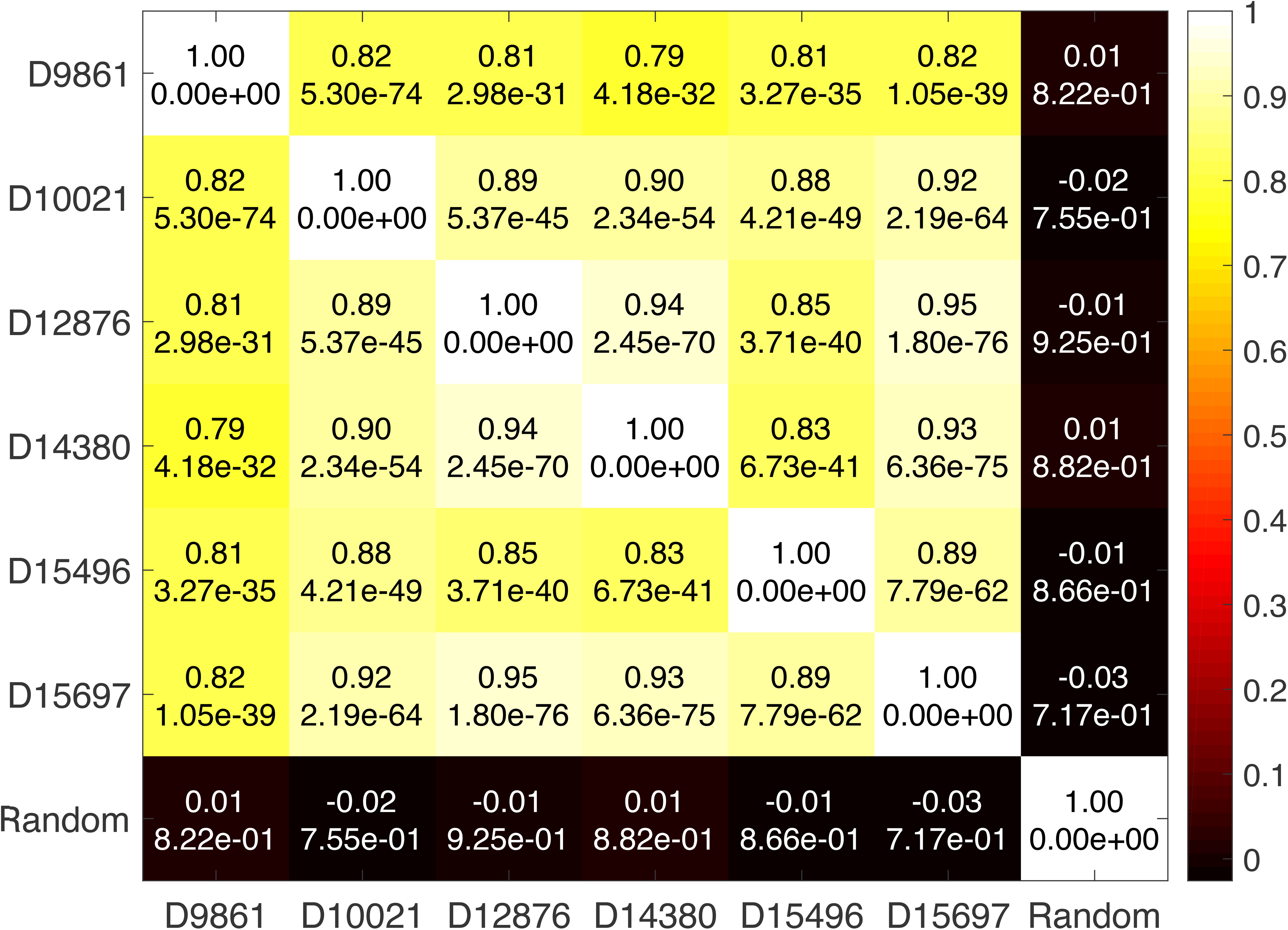
Allen Human Brain Atlas (AHBA) donor-to-donor consistency across MusNG, HumN and HumNG using the average Pearson’s correlation coefficient (PCC) of anatomic location and the p-value calculated from the average PCC.

### Cell-type distribution shows region specificity regardless of input data species

We observed that cell types with known anatomic locations mapped to the expected anatomic locations regardless of species. For example, the MusNG S1 cortex pyramidal cell type localized to the human cortex and the MusNG CA1 hippocampus cell type localized to the human hippocampus region in the AHBA donors (**Figure 5C2-3**, Supplementary Figure S14-31C2-3). Endothelial cell types also localized to human brainstem regions, where many large blood vessels enter the brain (Figure 5C6, Supplementary Figure S14-31C6). Ependymal cells localized to human ventricular regions of the brain (Figure 5C8, Supplementary Figure S14-31C8).

**Figure 5.**
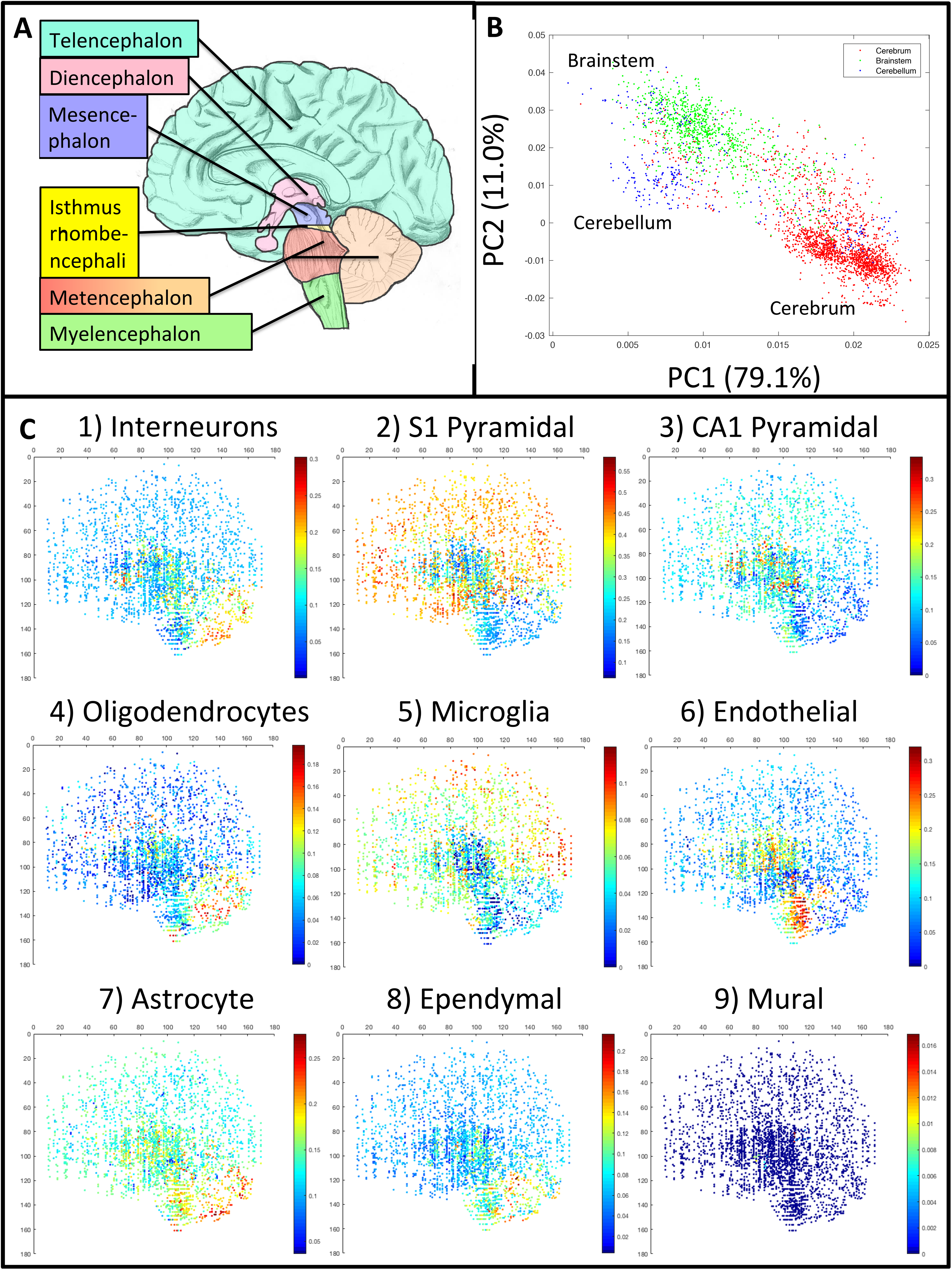
Spatial distribution of cell-type proportions in Allen Human Brain Atlas (AHBA) donors using all scRNA-Seq datasets to deconvolute the RNA expression profile. **A.** Reference map with developmental structures and major brain regions labeled. **B.** Principal component analysis (PCA) plot of the cell-type proportion matrix such that the nine cell types are reduced to two principal cell types for each of the samples in the brain. The colors indicate the three regions (cerebrum, brainstem, and cerebellum). **C.** The proportion of each cell type plotted from a sagittal view (see individual scale bars for proportions).

Brain localization information was known for certain cell types in each sample of the AHBA metadata files (points in Figure 5B,C). Clusters that emerged from principal component analyses (PCA) and K-means clustering were consistent with the sample’s brain region information. The average accuracy between the clusters and the true brain regions for all combinations of AHBA donors and scRNA-Seq datasets was 81 ± 7% (Supplemental Table I). The combined and smoothed example is shown in Figure 5B. The sensitivity and specificity of these clusters compared to the true brain region were also analyzed relative to the AHBA donor, scRNA-Seq dataset, and brain region (Supplementary Table II). We found high average sensitivity (0.80±0.16) and high specificity (0.91±0.05) across all scRNA-Seq datasets, brain donors, and brain regions (Supplementary Table II). The high sensitivity and specificity demonstrate that we can confidently identify the major brain region from which each sample is extracted using the cell-type proportions identified from our workflow. Supplementary Table II reports the sensitivity and specificity values for every combination of scRNA-Seq, AHBA donor, and brain region.

The multivariate analysis of variance (MANOVA) p-values for the sensitivity and specificity grouped by scRNA-Seq dataset, AHBA donor, and brain region were 0.3083, 0.2601, and 0.0019, respectively (Supplementary Table II, Figure 5B). These results showed that only brain region significantly affects the ability to identify the anatomical region from the cell-type composition. For example, it was more likely for brainstem samples to have cell-type compositions similar to the cerebellum or cerebrum than for cerebrum or cerebellum samples to have non-region-specific cell-type composition (Figure 5B, Supplementary Figure S14-31B, Supplementary Table II). More importantly, results were consistent for different scRNA-Seq input data or AHBA donor deconvolution. This finding further validates our feature selection and deconvolution techniques when mouse scRNA-Seq data is used to fill in human brain expression data for cell-type location analysis.

### Neuron nuclei and neuron mRNA localization are inconsistent when projection volume is not adjusted

The location of cells in a tissue is often considered to be as straightforward as the location of their nuclei. Using hematoxylin and eosin (H&E) staining, it is relatively straightforward to determine the locations of cell nuclei. In fact, nuclei stains are used to determine cellular location in applications like image segmentation [32, 33]. Consequently, cellular mRNA is considered in practice to follow similar spatial patterns to that of cellular nuclei. However, in cell types that may cover large distances, e.g., neurons, it may not be optimal to use nuclei location or mRNA location alone. We suggest that a measure of cellular extension, such as neural cell connectivity, should be used to better control for the mRNA contents of the cell with respect to the nuclei.

We found that the neuron/non-neuron ratios derived from mRNA deconvolution did not match those derived from counts of neuron nuclei [34]. The correlation between the nuclei-based ratios and our mRNA-based ratios across the cerebrum, brainstem, and cerebellum were not significant (**Figure 6A**; PCC = 0.1282, p-value = 0.3556). However, we noticed that the mRNA-based neuron/non-neuron ratios (p-value < 1.00×10^-16^ by ANOVA) and connectivity volume (p-value = 4.23×10^-15^ by ANOVA) varied significantly by brain region. When we divide the mRNA-based neuron/non-neuron ratios in each region by the mean projection volume for that region, the PCC significantly improves (Figure 6B; PCC = 0.7429, p-value = 1.26×10^-10^). We conclude that for neuron cell types, which have axons that travel long distances, the nuclei are not the optimal indicator for cell location. Instead, mRNA content information can better trace the cell location in brain tissue.

**Figure 6.**
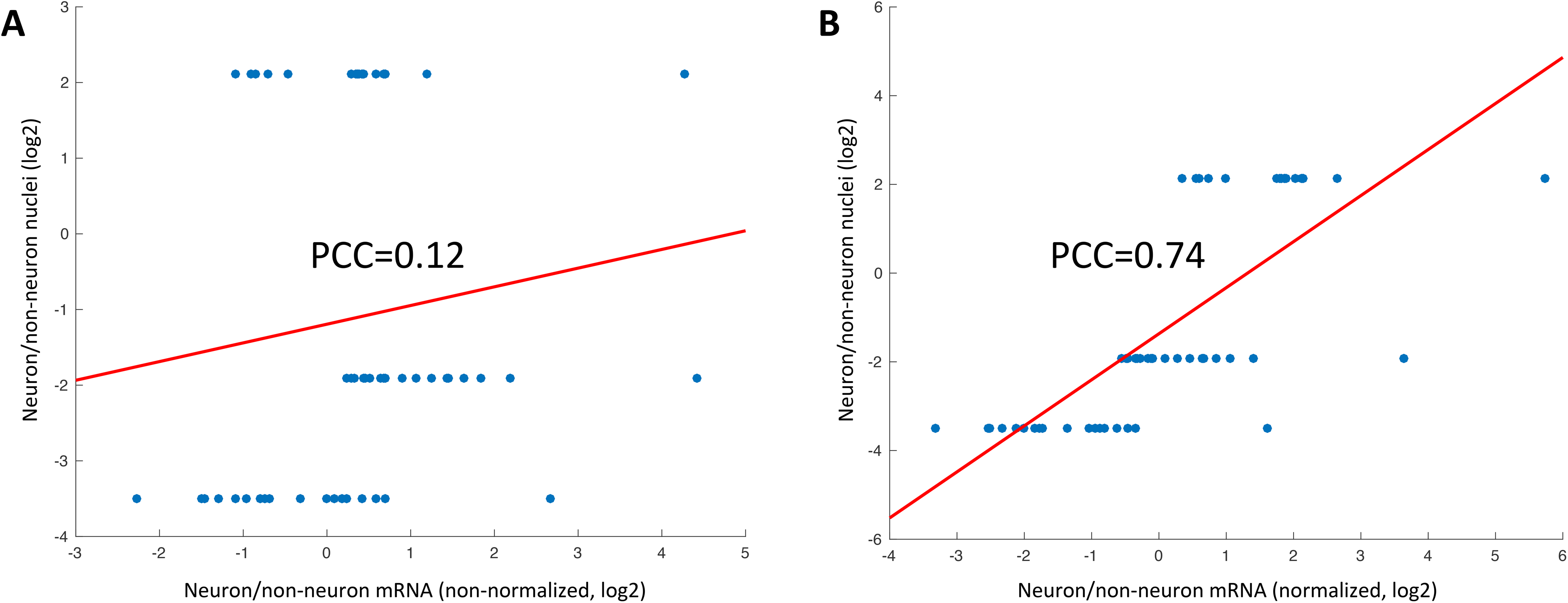
Comparison of nuclei/non-nuclei ratios derived from nuclei counts and mRNA quantification with respect to neural connectivity. **A.** The Pearson’s correlation coefficient (PCC) of the neuron/non-neuron ratios between the nuclei-based estimate and mRNA-based estimate. The 18 points along each tier of the y axis are the 3 RNA-Seq datasets by 6 Allen Human Brain Atlas (AHBA) donors. The three levels of nuclei ratios on the y axis are from the nuclei counts for each region. **B.** The data are the same as in A except that the mRNA neuron/non-neuron ratios were divided by the projection volume, which adjusts for the differences in neuron length among different regions.

### Principal cell types visualized across the entire AHBA

To visualize the structural patterns among estimated cell types, we applied singular value decomposition (SVD) to estimated cell-type data from the AHBA to reduce each mapped cell type to three principal types, which then could be displayed in an ℝ^3^ color vector. The 3D output for each of the six brains was overlaid by anatomic location. This visualization showed that there were unique patterns associated with cerebrum, brainstem, and cerebellum brain regions (**Figure 7A-I**). For example, the cerebrum displays a cell-type pattern that is distinct from that of both the brainstem and cerebellum. In contrast, although the brainstem and cerebellum exhibited differences from one another, the cell types within these two regions were similar (Figure 7A-I). These patterns were consistent across brains and among the input scRNA-Seq datasets that were used to deconvolute the samples (Figure 7A-I).

**Figure 7.**
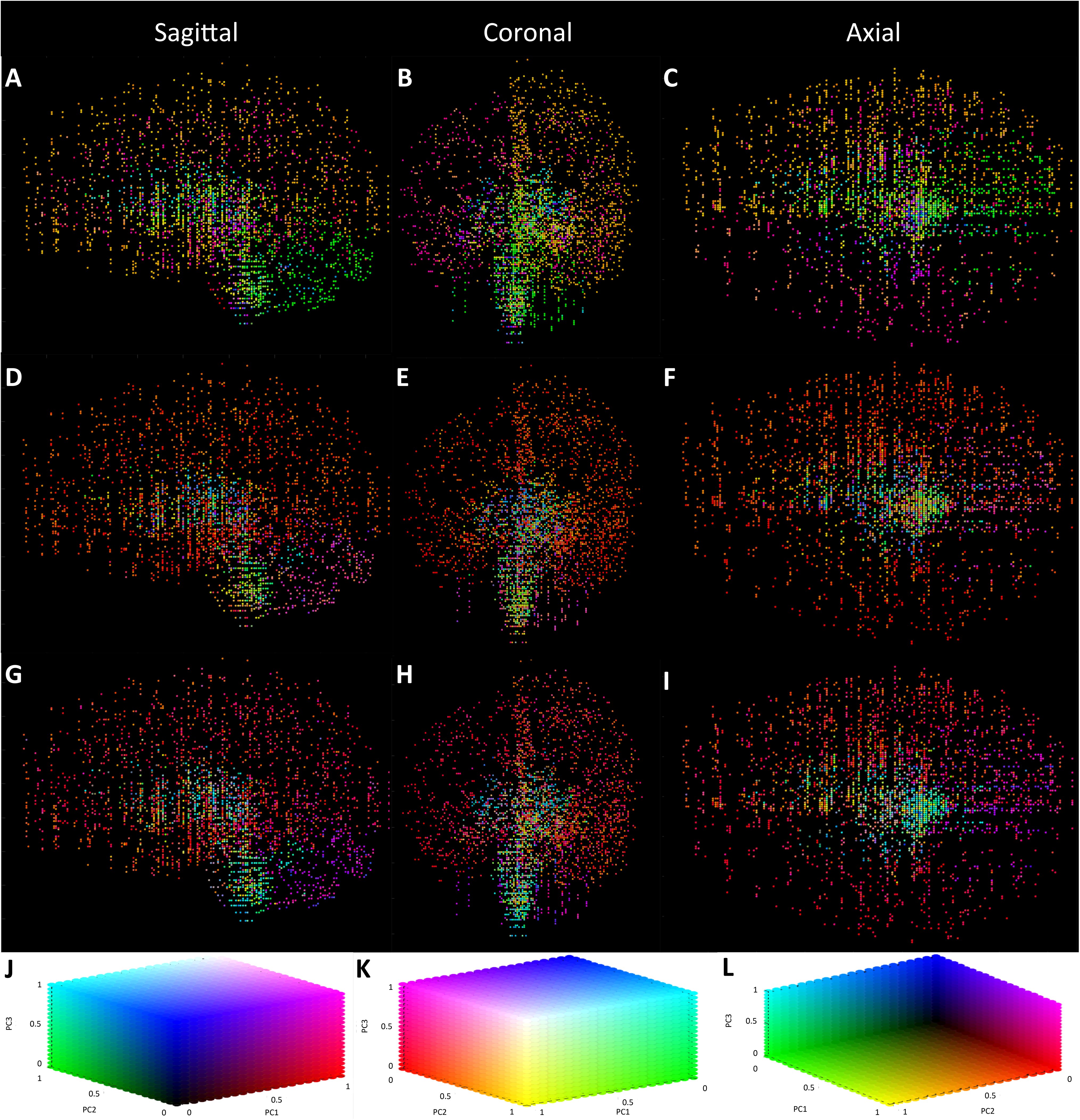
The 3D spatial mapping of major cell types from Allen Human Brain Atlas (AHBA) donors. Six samples are stacked together. The colors represent the first three principal cell types derived from principal component analysis (PCA) on the cell-type proportion matrix. **A, B, C**. MusNG deconvolution; **D, E, F**. HumN deconvolution; and **G, H, I**. HumNG deconvolution.

We also evaluated the principal cell types in each major brain region visually (Figure 7J-L, Supplementary Table III). The brainstem was generally comprised of CA1 pyramidal and glial cell types. The cerebellum was comprised of interneurons and various glial cell types (Figure 7J-L, Supplementary Table III). Though detailed information on specific cell-type locations are not common, it is worth noting that some patterns are consistent with known locations; for example, ependymal cells localized to the spinal cord and ventricular regions.

## Discussion

We demonstrated that brain cell-type density changes throughout the brain. We also showed that spatial information is consistent between species. Our methods successfully showed that cortex and hippocampal pyramidal cells in mouse also localized to cortex and hippocampus, respectively, in human samples. Furthermore, we showed that the neural cell-type proportions were generally much higher than the glial cell-type proportions. This finding may be due to a higher amount of total RNA in the cell or a greater diversity of RNAs. Alternatively, however, it is possible that neuron mRNAs were more specific to neurons, allowing the deconvolution algorithm to attribute a higher percentage of the tissue mRNA density to neurons. These considerations and their impact on spatial deconvolution could be partially accounted for by integrating nuclei density and neural connectivity data.

A previous study [34] focused on the cell-type composition of human brain tissue in specific anatomic brain regions by DAPI and NeuN protein staining in sections of human brain to count the number of nuclei in each section. The counts of all nuclei (DAPI+) and neuron nuclei (NeuN+) were used to extrapolate the proportions of neurons in a specific anatomic region. One striking finding from our study, however, is that regional neuron to non-neuron nuclei ratios did not accurately reflect cell-type specific mRNA composition when brain connectivity was not taken into account. This discrepancy indicates that nuclei counts are not necessarily adequate for evaluating the amount of neuron mRNA; for example, there may be more neurons in the cerebellum, but those neurons produce less mRNA than other brain regions. In addition, neural connectivity was greater in the cerebrum than in the cerebellum, despite greater numbers of cell nuclei in the cerebellum. These structural differences can be difficult to characterize at the single-cell level because the dissociation of cells in neural tissue severs these neural connections. For these reasons, therefore, we believe that more complex models should be developed to study cell-type information spatially in the brain. These models should take into account cell shape; changes in cell-type expression profiles by location; quantity of RNA in a given cell; and input from other data types, such as histological images. In addition to being integrative, these deconvolution models could also be more sophisticated by using machine learning approaches and feature reduction instead of feature selection.

Computational deconvolution to differentiate cell types is an important topic of transcriptomic data analysis, which can facilitate the elucidation of cell-type specific transcriptomic profiles in future studies [10, 19, 21, 23-31]. These approaches are generally limited to a single platform and usually for a single species, however, even though RNA-Seq/microarray data integration has already been applied in comparison studies [35, 36], tool development [37], and cancer research [38]. Due to the limitations of prior techniques, the vast accumulation of gene expression microarray and RNA-Seq data usually contains a tissue mixture with multiple cell types. With the single-cell RNA-Seq paradigm shift, it will be an important application of single-cell gene expression to deconvolute and “purify” this old microarray data to obtain cleaner representations of gene expression profiles for each cell type. In this study, we demonstrated that cell-type proportions could be derived with high accuracy through an integrated deconvolution method. We also achieved high consistency among scRNA-Seq datasets when they were used to deconvolute AHBA microarray data. We advocate for the improvement of this general methodology using integrative transfer-learning approaches to further evaluate both new and old datasets and to address problems associated with limited numbers of human brain donors.

The scarcity of human brain samples is a challenge in human neurological research. In most brain research, animal models are used as substitutes for human subjects. It is therefore important to integrate gene expression information from other species with human data. In fact, mouse brain expression can be used directly to study autism in the developing human brain [39]. Another study has shown that regional gene expression in mouse and human brain are concordant [18]. It has been shown that major neural cell types can be mapped between mouse and human if features are selected properly [17]. In this study, we found that we could achieve consistent cell-type deconvolution results in human brain samples by combining mouse and human data to deconvolute human tissue. We believe that this direct translation of mouse data to complement human data on homologous genes is a promising direction for transcriptomic and brain research. The additional data scope and sample availability from cross-species workflows is expected to facilitate the study of the structural, anatomic, and cellular heterogeneity that drives neurological development, cognition, disease, and degeneration.

## Conclusion

In this study, we estimated and visualized spatial cell-type changes across the entire AHBA by deconvoluting each of the 3,702 AHBA microarray samples with gene expression profiles of each cell type from both mouse and human scRNA-Seq. Highly consistent cell-type location patterns were achieved across all AHBA donors, which confirmed known cell types and location information. Furthermore, we found that the most conspicuous changes in cell types occurred within major anatomic regions, including the cerebrum, brainstem, and cerebellum. We also discovered unique spatial cell-type relationships, such as mouse hippocampal pyramidal cells localizing to human hippocampus. We also showed that both nuclei location and mRNA location should be considered when localizing neural cells, due to their large irregular shape. Furthermore, we advocate for the expansion of these proposed techniques to include more diverse data types and the integration of these techniques with more sophisticated transfer-learning approaches to generate more accurate models of tissue heterogeneity.

## Materials and methods

### Data

Ten datasets were used in our analysis. These datasets fall into three categories: 1) one mouse and two human scRNA-Seq datasets containing annotated cell types, which are used for expression deconvolution and cell-type specific signature generation; 2) six human microarray datasets from the Allen Brain Institute (AHBA datasets), which are used to generate human brain spatial maps; and 3) one mouse dataset of brain connectivity from the Allen Brain Institute, which is used to study the relationship between connectivity, cellular nuclei location, and cellular mRNA location in this work.

The mouse scRNA-Seq dataset [2] contains 3,005 cells, including both neural and glial cell types from the mouse hippocampus and cortex (denoted as MusNG, i.e., Mouse Neural-Glial). The two human scRNA-Seq datasets consist of different cell types. One dataset contains 3,086 cells (neural cell types only) from the cortex (denoted as HumN, i.e., Human Neural) [6]. The second dataset contains 285 cells (both neural and glial cell types) from the temporal lobe [5] (denoted as HumNG, i.e., Human Neural-Glial). Because MusNG is the most complete dataset in terms of the number of regions sampled and the diversity of cell types included, the MusNG dataset has been used to fill in missing cell-type expression data in the human datasets (Supplemental Figure S1).

The six microarray datasets that we used consisted of 3,702 total microarrays across every major brain region, and represent the entire set of expression data from the AHBA. Each microarray sample (tissue of mixed cell types) is taken for analysis from a specific location in one of the six donor brains. The 3D spatial coordinates of each microarray sample is also known. This information can therefore be used to map each microarray sample in 3D space. The datasets also include the brain region from which each sample is taken. A detailed description of how the data is generated and normalized can be found on the Allen Brain Institute website (http://human.brain-map.org/)

The brain connectivity dataset is generated from multiple mouse line brain tissue and contains all major anatomic regions (http://connectivity.brain-map.org). The sum of the projection volumes of both the injection and target sites are used as the measure of connectivity.

### Data preprocessing and feature selection

All mouse scRNA-Seq and AHBA microarray genes/probes are filtered prior to the analysis. Because the human scRNA-Seq data are missing several main cell types that are present in the mouse datasets, the mouse cell types are added to the human datasets to complete the human dataset. These appended mouse cell types are then subjected to downstream data filtering to prevent biases from being introduced. Rigorous data filtering is an important part of our analysis; scRNA-Seq data is sparse due to the nature of the lab preparation required to generate the reads as well as the cross-species/cross-platform nature of the data. The complete filtering takes four steps: 1) selection of mouse scRNA-Seq genes using the noise model [2]; 2) feature selection on mouse scRNA-Seq genes using minimum redundancy maximum relevance (mRMR) to remove highly correlated genes; 3) matching for homologs between mouse scRNA-Seq and human microarray probes; and 4) selection of concordant human and mouse gene homologs from the paired scRNA-Seq and microarray datasets. The details of each step are discussed in the preprocessing and feature selection section of the supplementary material (Supplementary Material, Section 1).

### Cell-type proportion estimates on simulated scRNA-Seq tissue

Ordinary Least Squares regression (OLS) and non-NMFR are two of the major deconvolution methods applied to gene expression data [22, 27, 40]. These two methods are tested in this paper for data integration and feature selection. To evaluate whether these methods are accurate in estimating the cell-type proportions, we apply both OLS and NMFR on simulated tissue samples generated from the scRNA-Seq datasets. These simulated tissue samples consist of aggregated single cells, which when combined represent a “simulated tissue” with known cell-type proportions (i.e., the counts of each type of cell in the aggregate). The accuracy of cell-type quantification is measured between the true cell-type proportions and the estimated proportions generated using OLS and NMFR. Based on the evaluation results, OLS is used to estimate the proportion of major cell types in the AHBA datasets. A detailed description of both regression methods can be found in the supplemental material (Supplementary Material, Section 2). For a gene expression data matrix E = ℝ^Gxc^, where each column constitutes one of c cell-type expression profiles with G gene features (rows), the proportions of each cell type as a vector (α_Tl_) can be estimated for a human brain sample *T*_*l*_, where *l* (1 ≤ l ≤ 6) represents the index of the sample in the total number of samples (*N* = 6) in the AHBA dataset.

### Cell-type proportion estimates across AHBA donors and scRNA-Seq datasets

Due to the complexity of cell types in brain tissues, it is necessary to employ multiple datasets (MusNG, HumN, and HumNG) to create the expression profiles used to deconvolute the human brain samples (*T*_*l*_). The consistency of these data types can be evaluated to ensure the method can be properly applied to human brain data. HumN contains only two major neural cell types, whereas HumNG contains one major neural cell type and four major glial types. MusNG is the most complete in terms of the major cell types (three neural and six glial/vascular types). Therefore, for any missing cell type in human scRNA-Seq data, it can be filled in using MusNG data such that the expression vector in MusNG corresponding to the missing vector in the human dataset is appended to the human dataset. Feature selection of concordant genes between species and sample-wise z-score normalization are conducted to make the vectors comparable. When comparing against HumNG, both MusNG and HumN aggregate S1 pyramidal and interneurons into a single neural cell type. For all datasets, CA1 pyramidal cells from MusNG is added because HumN and HumNG contain no CA1 pyramidal cells. A detailed account of how cell types are exchanged between datasets is provided in Supplementary Figure 1.

We employ the following methods to show that cell-type spatial maps in AHBA brains are consistent using deconvoluted signatures from any of the three scRNA-Seq datasets. For each dataset (MusNG, HumN, or HumNG), cell-type proportions are calculated across all six brains *[A^(s)(d)^]*. Let A^(s)(d)^ ∈ ℝ^Nxc^ s.t. 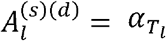 for scRNA-Seq dataset (s) and AHBA donor (d) and y^d^ the anatomic locations for each *T*_*l*_ in donor d. The computed cell proportions are subsequently used to calculate correlations between each pair of deconvolution input datasets. For each pair of comparisons, the mean correlation for each cell type from the six brains is used, resulting in 27 total correlations from three comparisons for nine cell types. We calculate the Benjamini-Hochberg false discovery rate (BH-FDR) for each of the 27 cell-type correlations. We also perform a Student’s t-test to compare the difference in PCC in the same cell types versus mismatched cell types.

It is also important to check the consistency between AHBA donors to ensure that our deconvolution results are valid. The A^(s)(d)^ are averaged across the three scRNA-Seq datasets (s), resulting in six averaged sets of proportions for each of the six AHBA donors. The brain regions for each sample are then used to create an average cell-type proportion for that region. The PCC is calculated between each of the AHBA donors using these regional proportions. A seventh random donor is created by i) randomly selecting a donor and ii) randomly reordering that donor’s sample regions. This donor is used to calculate a PCC between the random donor and the non-randomized donors. This randomization and correlation process is repeated 100 times and averaged to generate random PCC values as a negative control.

### Cell-type association with specific brain regions

To show the cell-type localization across all brains and across species/datasets, all scRNA-Seq datasets are used to deconvolute all AHBA brains using OLS. The results yield the sagittal view of the distribution of each cell type individually. The data from each scRNA-Seq dataset and AHBA donor combination are included and also combined into a comprehensive model. Each of the six brains are manually registered to each other so that the brain regions are consistent. Because some regions and cell types may have low representation in some samples, the deconvolution results (proportions) are smoothed by taking the maximum of the five closest samples in 3D Euclidean space. These new proportions are then returned to proportions such that each sample’s proportions sum to one, thus improving the coverage of difficult-to-detect cell types. The color represents the smoothed proportion of each cell type and is mapped to each sample’s respective voxel location in 3D space. To show overarching patterns, the principal cell types for that brain are plotted using a PCA plot. Each point (sample) in the PCA is colored based on the anatomic locations from which that sample was extracted in the donor brain. The PCs are calculated across the cell-type proportion matrix (A^(s)(d)^) for each scRNA-Seq dataset (s) and brain donor (d) using SVD, producing a matrix containing the PC values for each sample that could be plotted in 2D space. K-means clustering (k = 3, corresponding to the three major brain regions―cerebrum, brainstem, cerebellum―under study is applied to cluster the samples into three groups. The consistency between each of the three clusters and the three anatomic regions is measured using sensitivity and specificity values. A MANOVA model is fit using the sensitivity and specificity as dependent variables and the scRNA-Seq dataset, AHBA donor, and region sampled as the independent variables to study the effects of each type of sample on the ability to accurately map back to the tissue of origin. Any significant results for scRNA-Seq dataset, AHBA donor, or brain region correspond to a difference in mapping accuracy due to that variable (i.e., bias).

### Neuron nuclei and mRNA localization discrepancy and adjustment for neural projection volume

The inconsistency between neuron nuclei location and neuron mRNA localization is evaluated against neural projection information. We calculate the PCC between the deconvolution results and the nuclei count results, and then compare the PCCs while controlling for neural projection information. To explore the possibility that the long and irregular neuron volume could account for these discrepancies, we download the mouse brain connectivity data from the Allen Brain Institute to compute the neural projection volume. First, the injection and target site volumes for each experiment as well as the mouse anatomic brain region hierarchy are downloaded. A breath-first-search is employed to extract all regions under the cerebrum, brainstem, and cerebellum branches in the mouse brain region hierarchy. This information is used to stratify the signals to the specified regions. Next, the metadata from each of the AHBA donor brains is used to extract cerebrum, brainstem, and cerebellum samples, stratifying the sample proportions to each region. The ratio of neuron to non-neuron signature is calculated for each sample and used in subsequent analysis so that the cell-type signature proportions are comparable to the nuclei proportions. Next, all 18 combinations (three input scRNA-Seq datasets and six brain donors) are matched to each of the nuclei datasets by brain region such that for each nuclei neuron/non-neuron estimate, there are 18 cell-type signature proportion-derived neuron/non-neuron estimates. Finally, the cell-type expression profile proportion ratios are divided by the axonal projection volumes (mouse connectivity) to recreate a positive correlation between nuclei and cell-type expression profile proportions.

### Visualization of principal cell types across the entire AHBA

To show high-level spatial distributions of principal cell types, all six brains across all cell types are combined into single 3D representations for each of the three scRNA-Seq datasets. SVD was performed on the cell-type proportion matrix, and the three largest components were used as the <u>principal cell types</u> across all samples in each brain to display the results as a three-digit color vector. This analysis is performed for each of the six AHBA brains using each of the three scRNA-Seq datasets individually. The principal cell types are displayed in 3D using color to represent the top three principal cell types. The three scRNA-Seq datasets are displayed separately such that all six AHBA brains are manually overlaid for each scRNA-Seq dataset. This registration produces consistent anatomic locations across the six overlaid brains. Displays are generated with MATLAB function scatter3 (The Mathworks, Inc.; Natick, MA, USA) to view sagittal, coronal, and axial images.

## Authors’ contributions

TSJ, ZBA, and PN performed data analyses. KH, RM, and TSJ conceived and initiated this project. JZ, YZ, and KH supervised the project. TSJ, ZBA, YZ, KH, and JZ contributed to biological interpretation. TSJ, ZBA, BRH, YZ, KH, and JZ wrote the manuscript. All authors read and approved the manuscript.

## Acknowledgments

This research was supported by a National Institutes of Health NLM-MIDAS Training Fellowship (4T15LM011270-05) to TSJ and The Ohio State University (Columbus, OH) and departmental start-up funding from the Indiana University School of Medicine (Indianapolis, IN) to KH.

