## Supplementary Materials for "Integration of Mouse and Human Single-cell RNA Sequencing Infers Spatial Cell-type Composition in Human Brains"

**Supplementary Information**

**Sec. 1 Cleaning and Feature Selection**

*Step 1: Noise Model Filtering by Selection of Genes and Clustering of Mouse Cells*

The raw unique molecular identifier (UMI) counts obtained from the MusNG dataset were cleaned and filtered in the same manner as Zeisel et al^12^. This process took three steps: i) removing low expression genes, ii) removing genes that do not co-express with other genes and iii) selection of top 5000 high variance genes. The intersection of the 5000 MusNG genes with the complete HumNG dataset resulted in 4009 genes that were used in the subsequent analysis. The HumN dataset was not reduced to these 5000 genes because there were too few cell types to perform the mRMR feature selection on this dataset alone.

*Step 2: Feature Selection of Genes in scRNA-Seq Data*

After filtering, an open-source feature selection algorithm was used to further reduce the number of redundant genes that coexpress within the cortex dataset. mRMR was used to select the top 500 genes from the original 5,000 in the MusNG dataset and 4009 in the HumNG dataset. This tool finds features that are “mutually as dissimilar to each other as possible, but marginally as similar to the classification variable as possible” ^36^. This method reduces the number of redundant features that would be included in the cell type models.

*Step 3: Matching homologs between Mouse scRNA-Seq and Human AHBA Microarray*

After the feature selection step the mRMR-selected genes were then paired with microarray data obtained from AHBA donors individually. The pairing further reduced the number of genes to a subset of genes that were contained in both AHBA and the scRNA-Seq datasets. MusNG contained 423 genes, HumN contained 382 genes, and HumNG contained 62 genes.

*Step 4: Selecting Concordant Homologs Between Mouse scRNA-Seq and Human AHBA Microarray*

These remaining genes were then center normalized (so that all cell types do not have a higher mean expression across all genes), log2 transformed and converted to a z score. For HumN and HumNG the missing cell types were filled in using the more complete MusNG dataset such that HumN contained 2 human and 7 mouse cell types and HumNG contained 5 human and 3 mouse cell types. Because the AHBA data is fundamentally different due to platform and species genes needed to be selected that behaved the same across the datasets. This introduced a problem in that a traditional correlation matrix method cannot be computed because the microarray and scRNA-Seq samples are not paired by independent variables. However, gene lists ranked by correct classification of brain regions correlate between mouse and human ^37^ and mouse and human genes can be selected that will correctly cluster cell types regardless of species^38^. Based on this evidence, we developed a rank based approach using the difference in z-scores of both log2 transformed microarray and scRNA-Seq data matrices to identify concordant homologs. An allowable difference in standard deviation (standard deviation constant) was chosen that corresponded to the maximum allowable change between the mean z score for each gene across all samples.

To attain this, a vector of mean expression (each element for one gene) across all microarray samples was used to compare against the scRNA-Seq data.

$\bar{T}= \frac{1}{N}\sum_{1}^{N} T_{l}$	(eq. 1)

Where $T_{l}$ is expression vector from sample $l$. Next a vector was generated from the RNA-Seq CTEPs – a weighted average of all CTEPs initialized to a uniform distribution ($\hat{E})$.

$\hat{E}=Eb^{(j)}$	(eq. 2)

Where $E= \mathbb{R}^{length(G)\times c}$ is the expression RNA-Seq expression matrix (columns are cell type profiles rows are genes) and $b^{(j)}$ contains the weights for each CTEP, $b= \mathbb{R}^{c} s.t. b_{i}=\frac{1}{c}\forall i\mathbb{\in Z,}1\leq i\leq c$. Variable $j$ represents the iteration of the algorithm such that when first initialized $j=0$. Then all genes were selected that are within a standard deviation constant of one another.

$kept\subseteq G s.t.\left| \bar{T}^{(g)}- \hat{E}^{(g)} \right| \leq d \forall g\in G$	(eq. 3)

Where $kept$ is the selected gene set, $T\in\mathbb{R}^{length(G)\times N}$ is the microarray expression matrix (columns are samples and rows are genes), $G$ is the set of all genes, $d$ is the standard deviation constant and $g$ is a gene that is within d units of itself between the RNA-Seq and microarray dataset. The predicted proportions ($\hat{b}$) of each CTEP in $\hat{T}$ were calculated using the retained set of concordant gene features ($kept$).

$min(\left| E^{kept}\hat{b}- \bar{T}^{kept} \right|)$	(eq. 4)

$b^{(j+1)}=z\hat{b}+(1-z)b^{(j)}$	(eq. 5)

Where $z\mathbb{\in R}s.t. 0 \leq z \leq1$ is the step size for each iteration of our feature selection algorithm (0.3 was used in this study due to the inflection point in Supplementary Figure 3)$.$Variable $b^{(j+1)}$ was then incremented with the updated predicted proportions of CTEPS ($\hat{b}$) (eq. 5) then $j$ was incremented ($j=j+1)$. The error was then calculated.

$error=\left| E^{kept}\hat{b}- \bar{T}^{kept} \right|$	(eq. 6)

These steps (2-5) were repeated until there was no longer a decrease in the error between steps.

The standard deviation constant ($d$) and step size ($z$) have unique features that can have effects on the estimates (Supplementary Figure 3). High standard deviation constants correspond to more genes that have a lower correlation between the datasets and low standard deviation constants correspond to fewer genes that have a higher correlation between the microarray and scRNA-Seq datasets. The standard deviation constant was set to 0.9 for most of the analysis and step size to 0.3 due to the inflection point shown in (Supplementary Figure 3).

**Sec. 2 Deconvolution Techniques**

### Ordinary Least Squares Regression (OLS)

The tissue level expression can be thought of as the weighted mean of the cell types within that sample ^25,31,42^. Eq 1, the linear combination of $c$ cell types within the tissue sample $T_{l}$, easily represents this.

$T_{l}=\sum_{i=1}^{c} E_{i}\propto_{i}+\epsilon$	(Eq 7.)

Where α_i_ is the proportion ($0 \leq\propto_{i}\leq1,\sum_{i=1}^{c} \propto_{i}$ = 1, $i\mathbb{\in Z},1\leq i\leq c$) of cell type $i$ in the tissue sample $T_{l}$, $E_{i}$ is the expression profile of cell type $i$ and $\epsilon$ is the error vector. This can be re-written as Eq 2.

$T_{l}=E\propto+\varepsilon, E\in\mathbb{R}^{kept\times c}, \propto\in\mathbb{R}^{c}$	(Eq 8.)

A MATLAB script was used to fit a linear model to each of the samples within each AHBA donor using each of the three scRNA-Seq dataset derived CTEPs. The MATLAB function lsqnonneg was used to calculate the α values from the mean expression levels across the retained genes that were selected for the cell type model in each of the tissue samples. The ordinary least squares regression (OLS) method does not include a feature reduction step, which in some applications could be detrimental (e.g., clustering). However, due to the filtering of redundant features in the dataset using mRMR much of the feature reduction had already been performed.

### Non-negative Matrix Factorization Regression (NMFR)

NMFR is based on Principal Components Regression (PCR), an alternative to OLS, which takes advantage of existing variance in the data by fitting new components to the data that maximize the data variance ^43^. In the case of PCR, a score matrix ($S$) and loading matrix ($P$) is calculated from $E$ (the matrix of concatenated $E_{i},i\mathbb{\in Z,}1\leq i\leq c$) such that $S_{j}$is the $j^{th}$ principle component of the model—the component with the $j^{th}$ highest variance. Regression can be run on the principle components (score matrix $S$).

$T_{l}=S\beta+\varepsilon, S\in\mathbb{R}^{kept\times r},\beta\in\mathbb{R}^{r}$	(Eq 9.)

Where $\beta$ is the weight of principle component j in the linear combination of principle components in $T_{l}$, while $S_{j}$ is the expression profile of principle component $j$. To retrieve the cell type specific proportions ($\propto$) from the PCR, $\beta$ can be transformed back into the original variables ($\propto$) using the loading matrix $P$.

$\propto=\beta^{T}P, P\in\mathbb{R}^{r\times kept}$	(Eq 10.)

However, the main problem encountered when using PCR to deconvolute cell types is that the proportion of each cell type is not by definition non-negative. To avoid this problem, non-negative matrix factorization (NMF) can be used instead to create a lower dimension matrix from the original cell type matrix with c columns, corresponding to each CTEP and kept rows corresponding to the selected features (genes). NMF is a technique where a matrix can be approximated into the product of two non-negative factor matrices ^44,45^. Let $E$ be the raw expression matrix.

$E\approx WH,E\in\mathbb{R}^{kept\times c},W\in\mathbb{R}^{kept\times r},H\in\mathbb{R}^{r\times c}$	(Eq 11.)

The non-negative matrix factorization of $E$ produces $W$ and $H$ which can be thought of as the non-negative score matrix and loading matrix respectively. By treating the non-negative approximation matrix $W$ as the principal components and the non-negative fit function used in the OLS model, non-negative $\beta$ values are produced that can then be used as estimates of cell type densities. Eq 9 and 10 can be rewritten in terms of $W$ and $H$ to attain non-negative estimates.

$T_{l}=W\beta+\varepsilon$	(Eq 12.)

$\propto=\beta^{T}H$	(Eq 13.)

**Sec. 3 Significance testing of cell type correlations**

To test whether the predicted cell type proportions were consistent in their spatial distribution regardless of the starting dataset (MusNG, HumN, HumNG), we calculated correlations and performed significance testing. As described in the main text we calculated the correlation of each cell type to itself across starting datasets. These correlations were used in the main text Figure 3. We also calculated every combination of cell types within dataset and across datasets for each starting dataset comparison. This resulted in a correlation matrix where all cell type correlations could be studied (Supplementary Figure 4-10). Since the overall cross dataset correlations are of interest we also display the mean correlation across all brains for each of the three comparisons (MusNG+HumN, MusNG+HumNG, HumN+HumNG, Supplementary Figure 4). We find that in this overall comparison the same cell type comparisons are always more positively correlated than mismatched cell type correlations (Supplementary Table 1).

**Supplementary Figures**

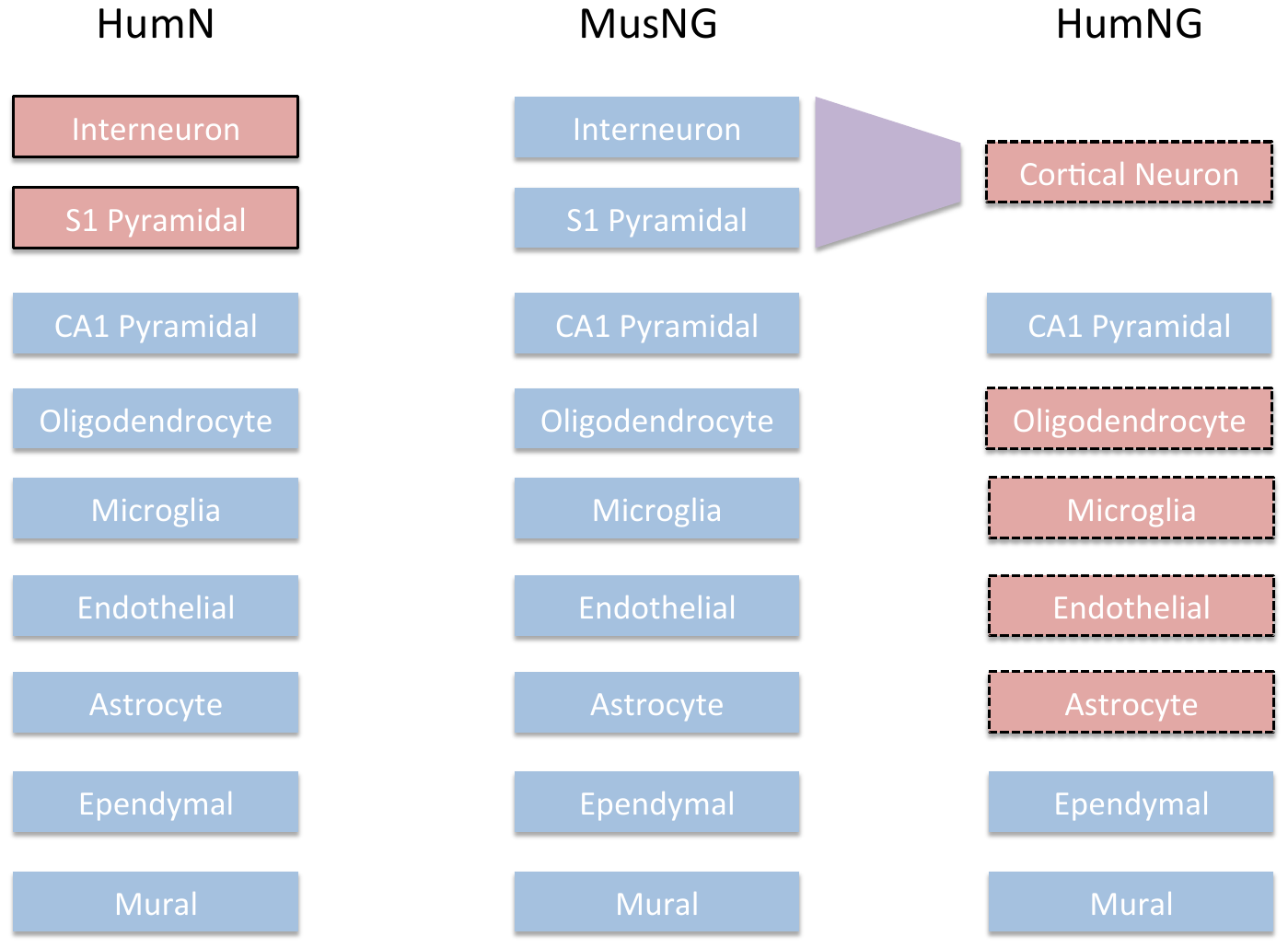

Supplementary Figure S1 To have comparable datasets some cell types had to be used from the mouse dataset to complete the two human datasets. Red boxes indicate human cell types and blue indicate mouse cell types. The original HumN cell types are outlined in a solid line. The original MusNG cell types have no outline. The original HumNG cell types are outlined in a dashed line.

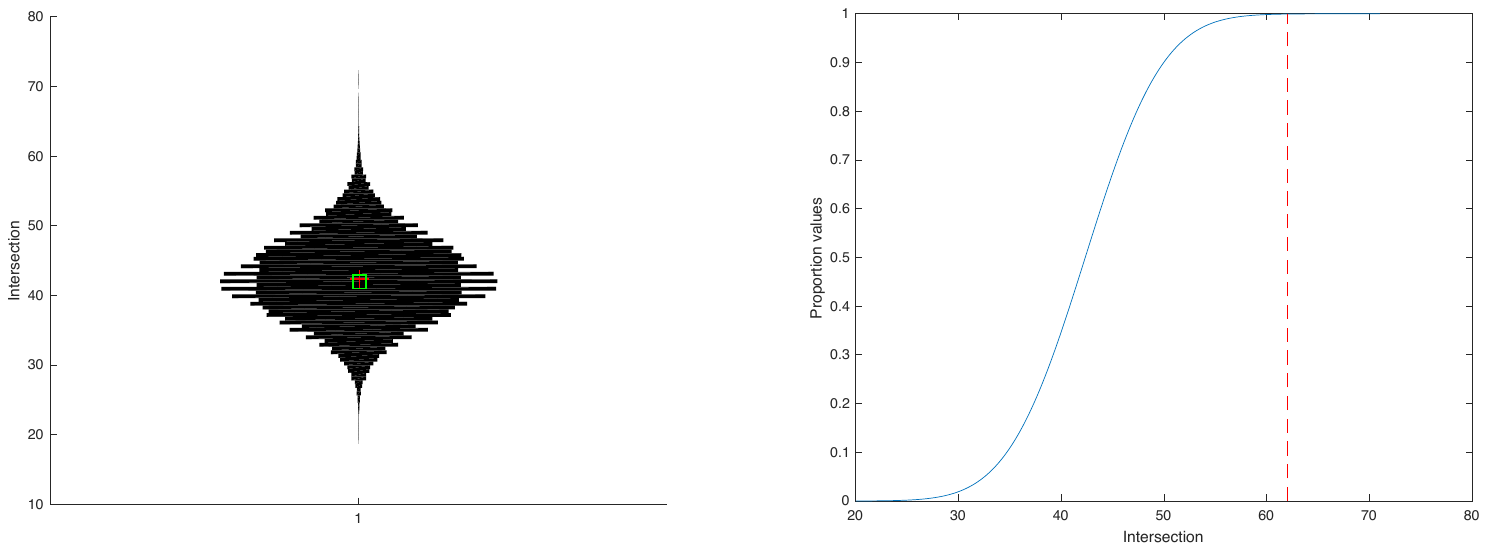

Supplementary Figure S2 This plot represents the empirical study of the likelihood of acquiring an intersection of 62 features from 2 randomly selected samples described in the discussion section. Left is the distribution of intersections between two randomly selected datasets. Right is the CDF of that distribution with the intersection found during our study marked by the red dashed line.

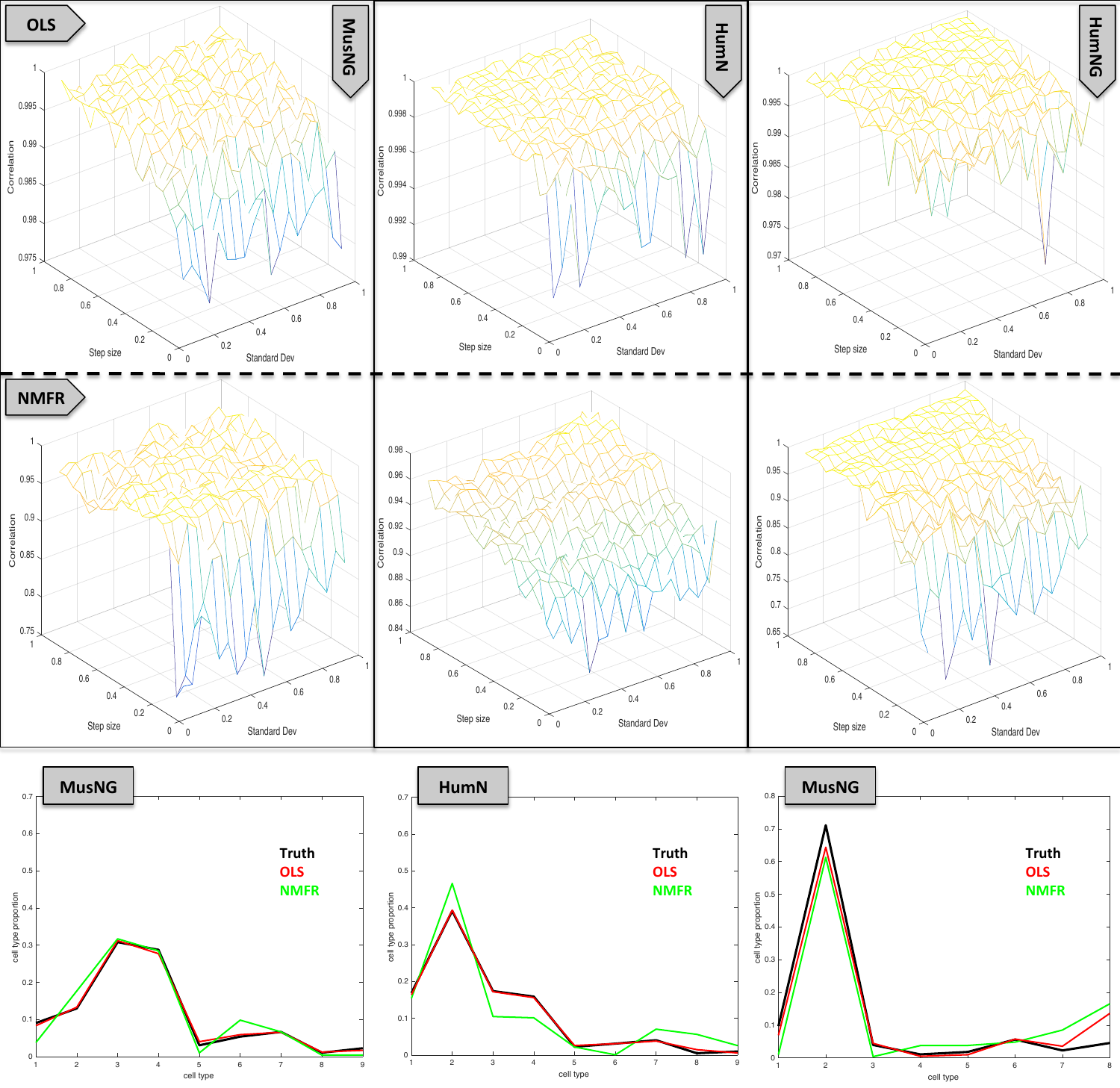

Supplementary Figure S3 Each plot represents the changes in correlation between the estimated and true cell type proportions with the change in step size (z Eq. 5) and (d Eq. 3). The rows represent the algorithm used to estimate the proportions and the columns represent the scRNA-Seq datasets used to generate the artificial sample and cell type expression profiles.

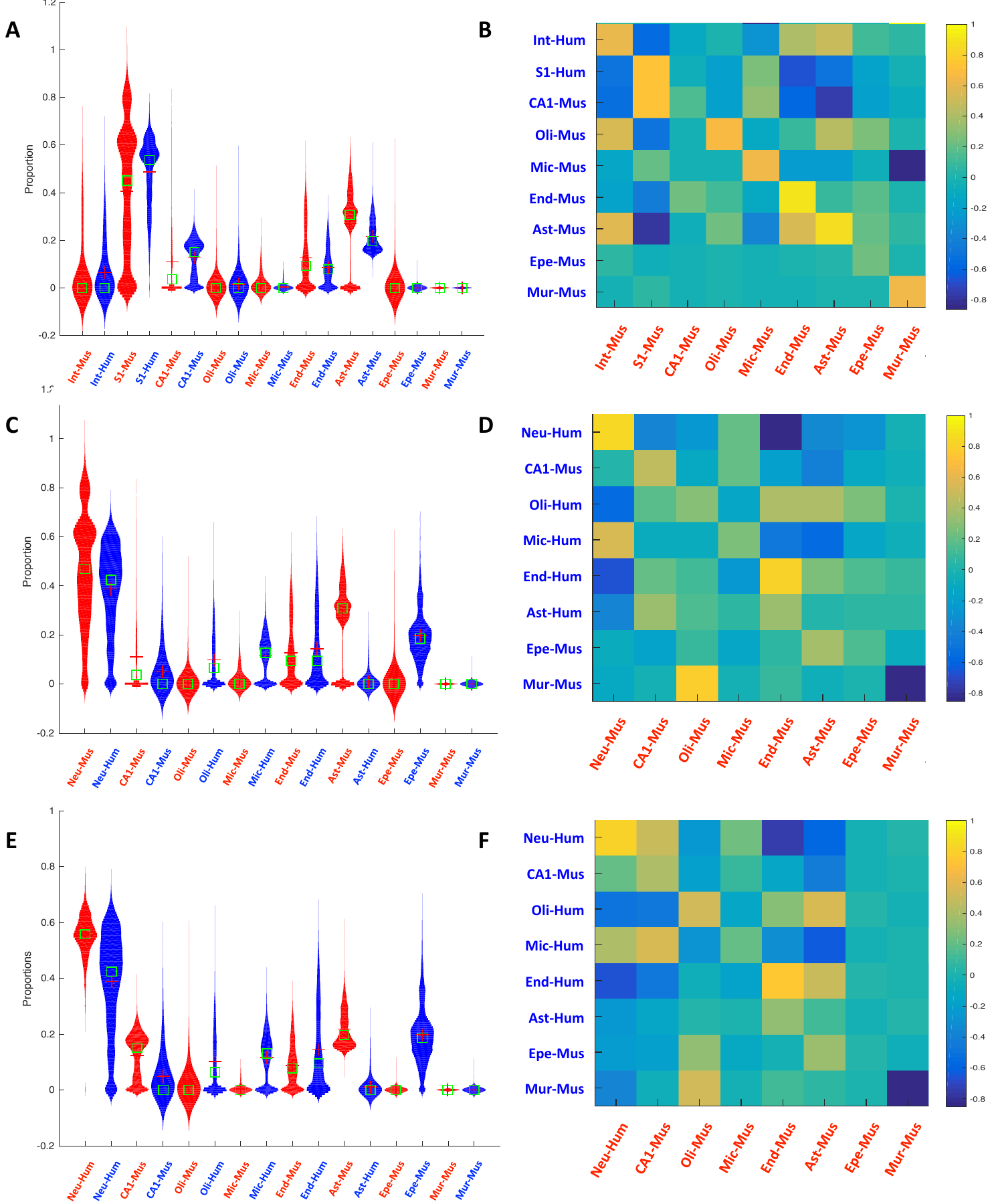

Supplementary Figure S4 Consistency of cell type proportions using each of the scRNA-Seq datasets to deconvolute the AHBA. Each figure is the combinations of all 6 AHBA donors. A) Violin plot of MusNG (red) and HumN (blue) distributions of cell type proportions for each cell type. B) The cell proportion correlations for each cell type originated from MusNG (blue) vs. from HumN (red) C) Violin plot of MusNG (red) and HumNG (blue) distributions of cell type proportions for each cell type. D) The cell proportion correlations for each cell type originated from MusNG and HumNG . E) Violin plot of HumN (red) and HumNG (blue) distributions of cell type proportions for each cell type. F) The cell proportion correlations for each cell type originated from HumN and HumNG . B,C,D) The red fonts correspond to the red distribution and the blue fonts to the blue distribution plots in A,C,E.

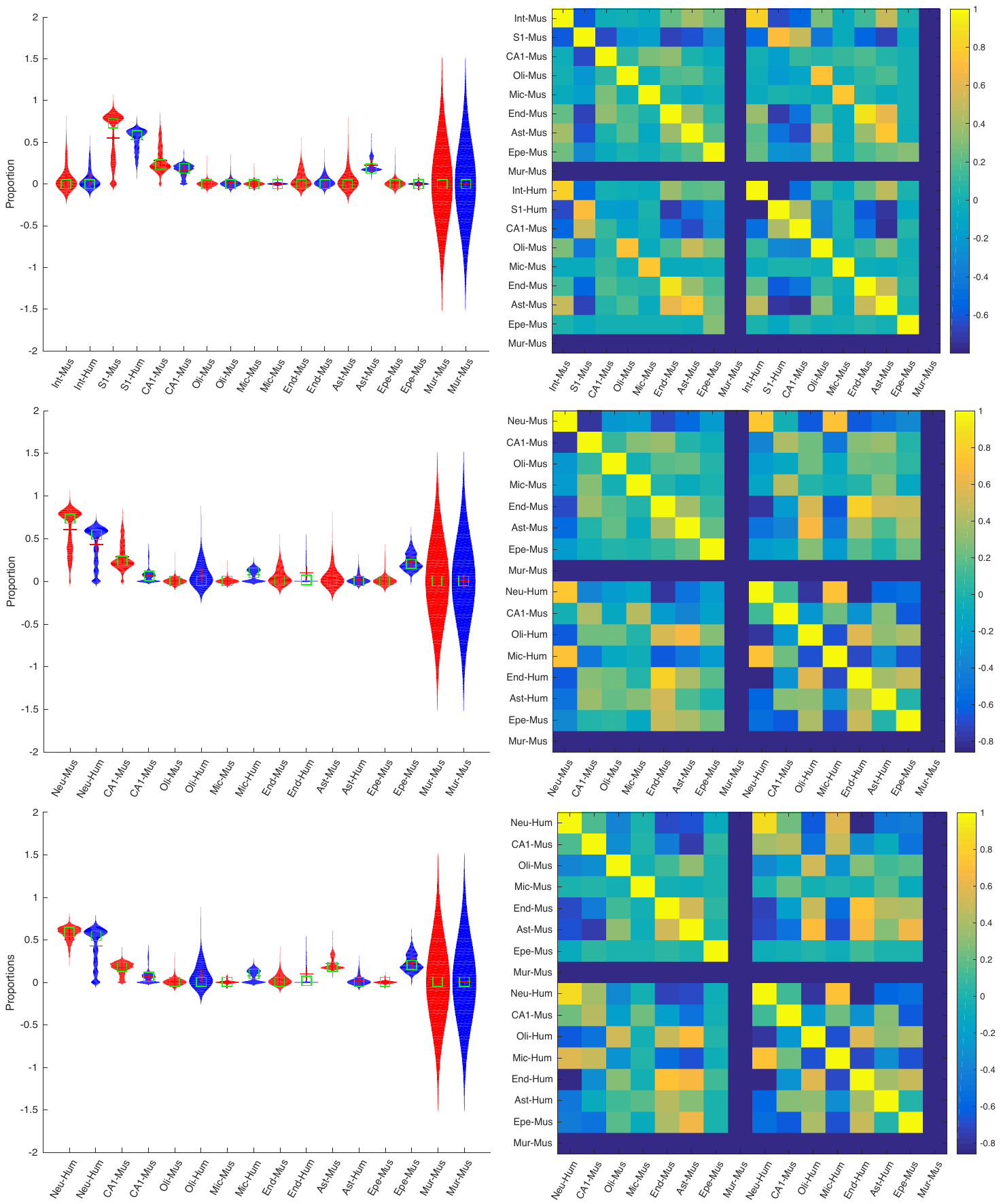

Supplementary Figure S5 Comparison of cell type distributions between each combination of datasets within donor 9861. Top: MusNG (red) and HumN (blue), Middle: MusNG (red) and HumNG (blue), Bottom: HumN (red) and HumNG (blue). Left column are the cell type distributions from donor 9861 such that each cell type signature distribution (x axis) contains every observed proportion for that cell type across all samples in donor 9861’s brain. Right column are the spatial correlations between cell type proportions between datasets (p-values in Supplementary Table 2).

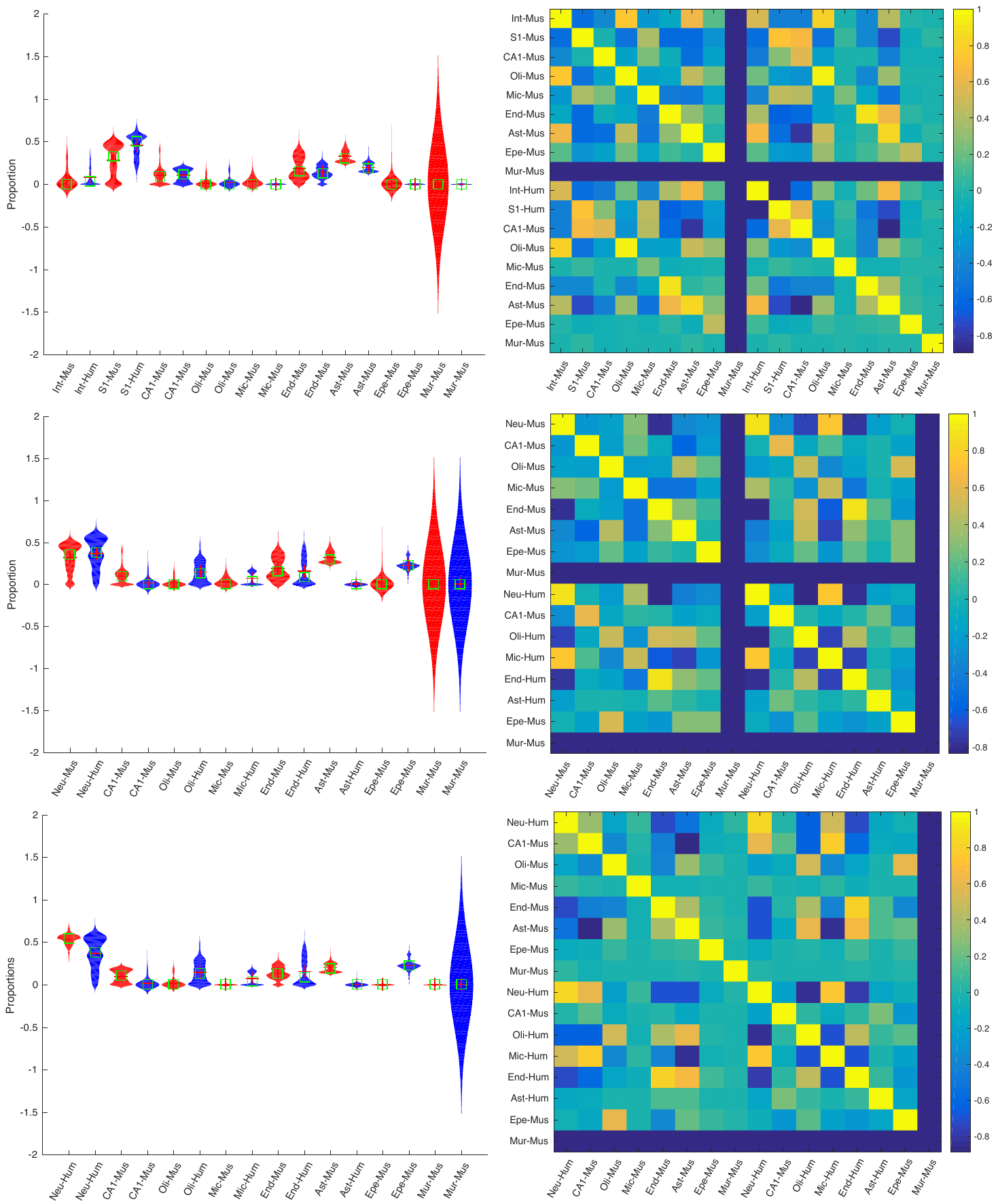

Supplementary Figure S6 Comparison of cell type distributions between each combination of datasets within donor 10021. Top: MusNG (red) and HumN (blue), Middle: MusNG (red) and HumNG (blue), Bottom: HumN (red) and HumNG (blue). Left column are the cell type distributions from donor 10021 such that each cell type signature distribution (x axis) contains every observed proportion for that cell type across all samples in donor 10021’s brain. Right column are the spatial correlations between cell type proportions between datasets (p-values in Supplementary Table 2).

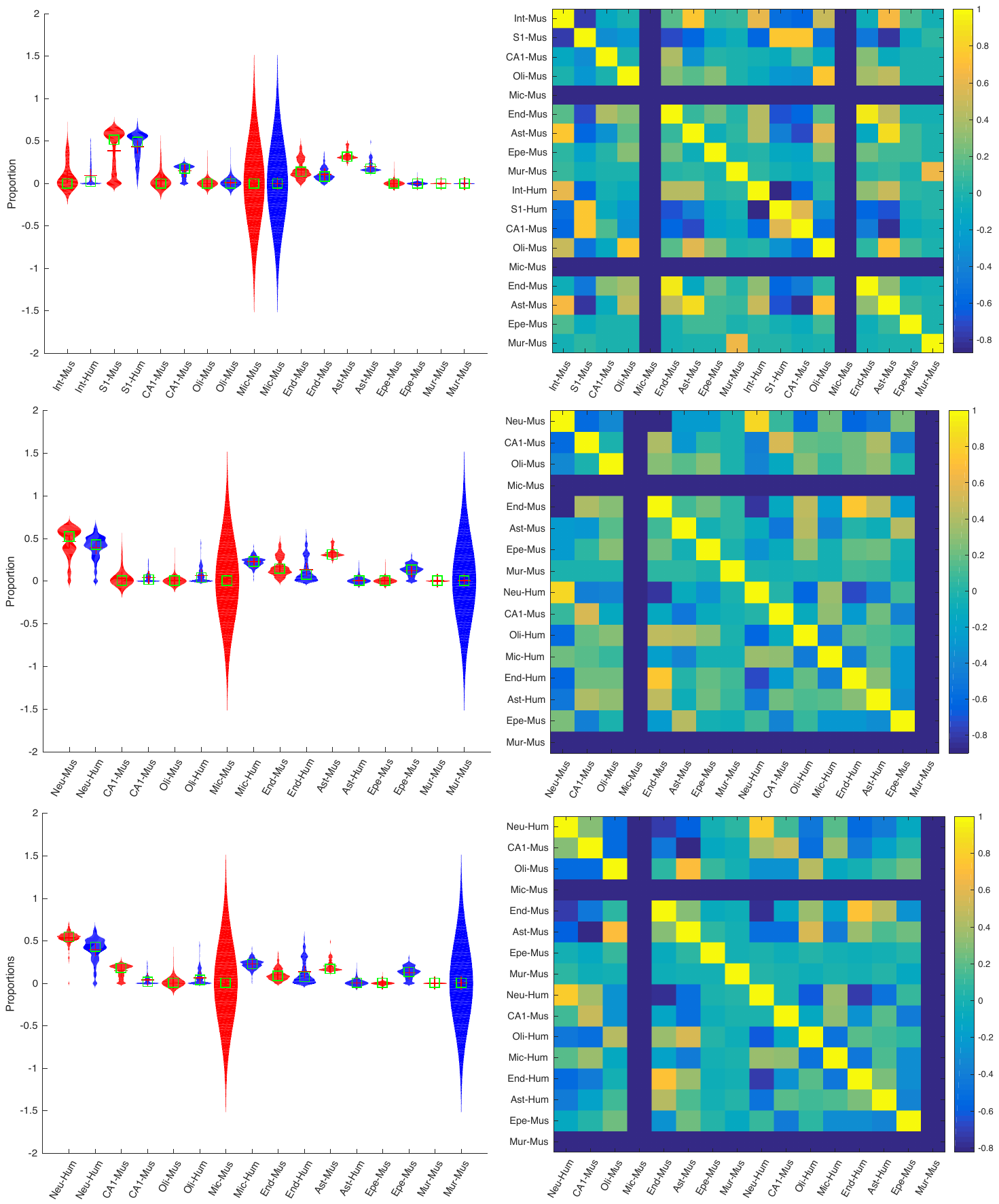

Supplementary Figure S7 Comparison of cell type distributions between each combination of datasets within donor 12876. Top: MusNG (red) and HumN (blue), Middle: MusNG (red) and HumNG (blue), Bottom: HumN (red) and HumNG (blue). Left column are the cell type distributions from donor 12876 such that each cell type signature distribution (x axis) contains every observed proportion for that cell type across all samples in donor 12876’s brain. Right column are the spatial correlations between cell type proportions between datasets(p-values in Supplementary Table 2).

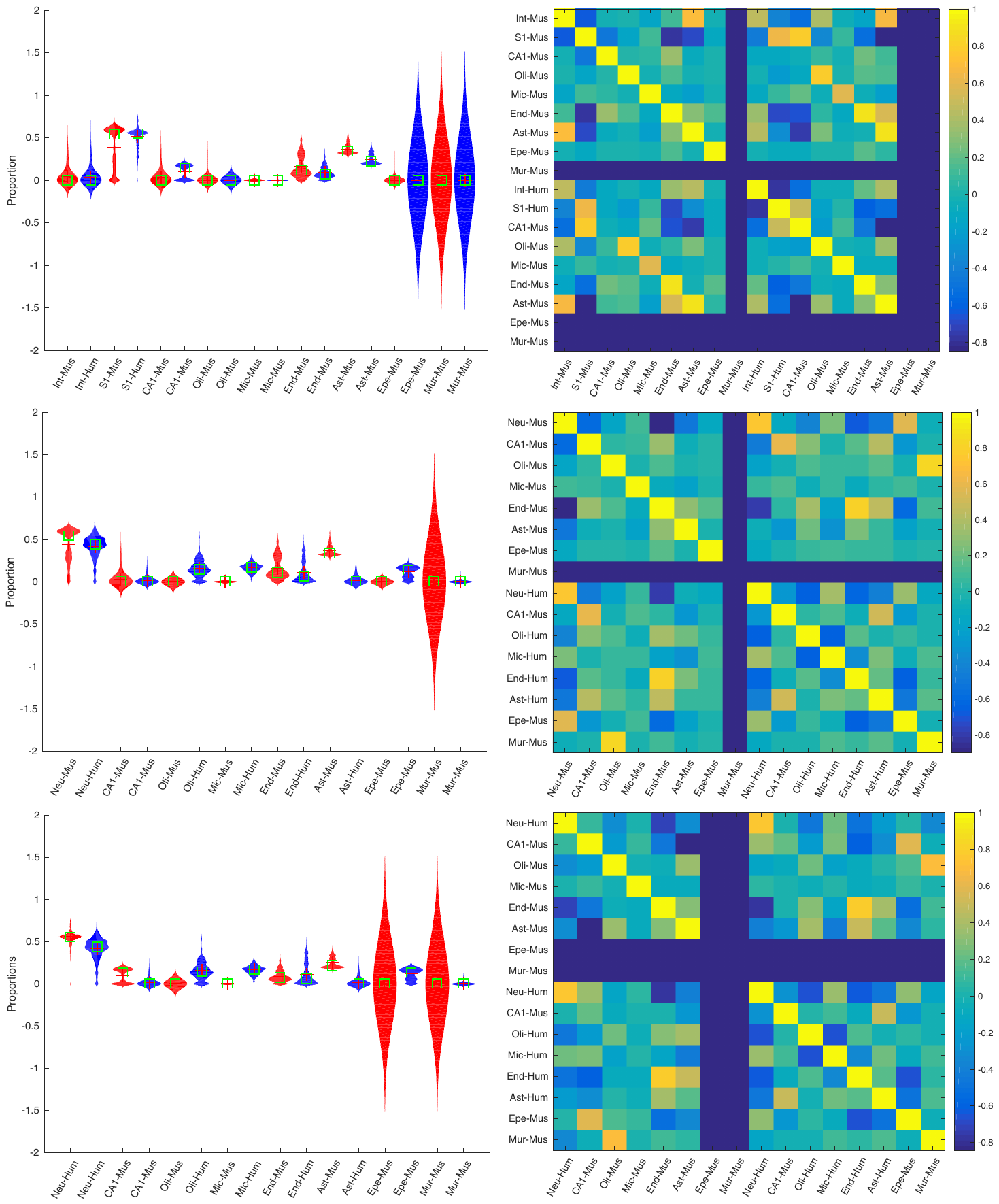

Supplementary Figure S8 Comparison of cell type distributions between each combination of datasets within donor 14380. Top: MusNG (red) and HumN (blue), Middle: MusNG (red) and HumNG (blue), Bottom: HumN (red) and HumNG (blue). Left column are the cell type distributions from donor 14380 such that each cell type signature distribution (x axis) contains every observed proportion for that cell type across all samples in donor 14380’s brain. Right column are the spatial correlations between cell type proportions between datasets (p-values in Supplementary Table 2).

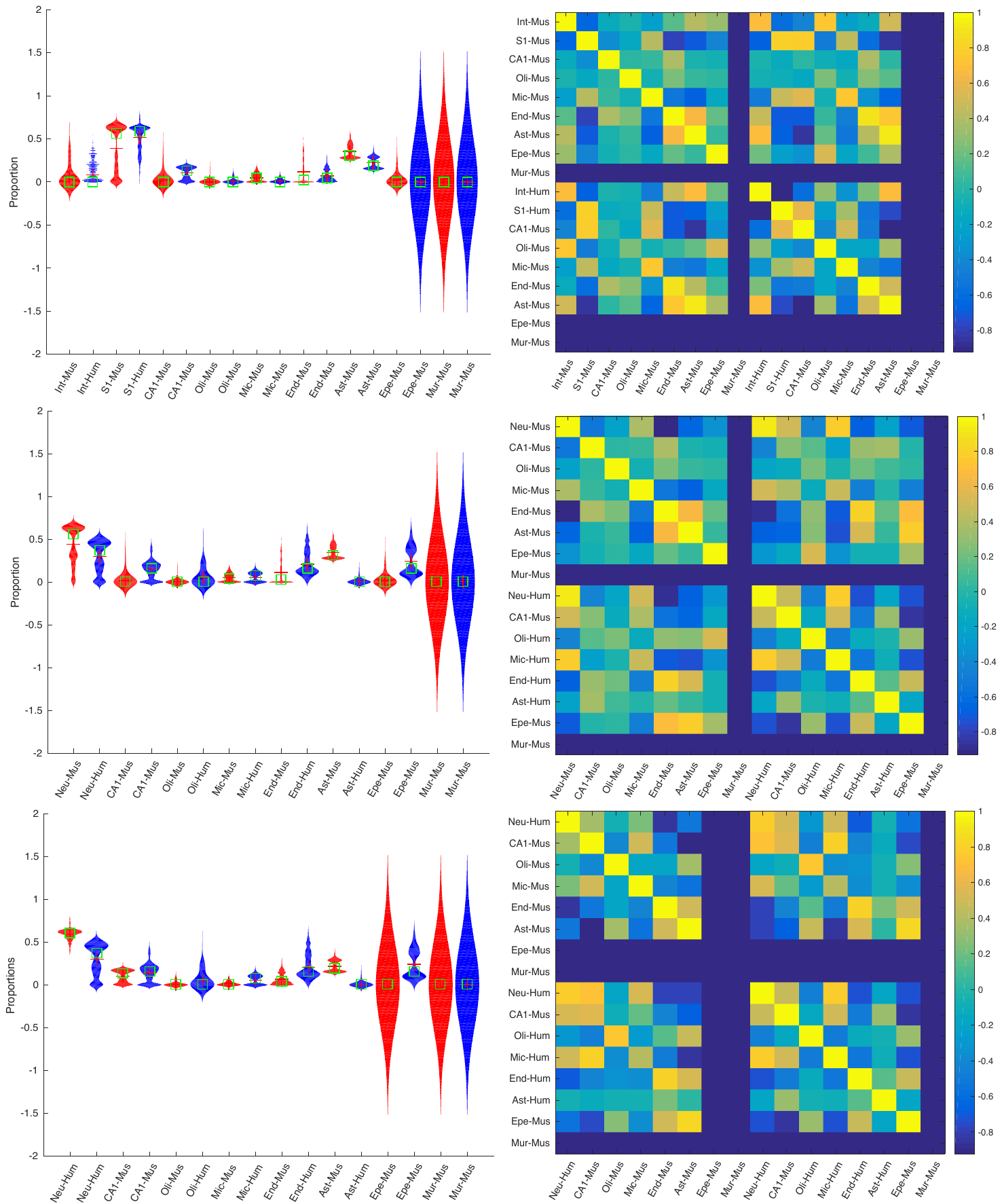

Supplementary Figure S9 Comparison of cell type distributions between each combination of datasets within donor 15496. Top: MusNG (red) and HumN (blue), Middle: MusNG (red) and HumNG (blue), Bottom: HumN (red) and HumNG (blue). Left column are the cell type distributions from donor 15496 such that each cell type signature distribution (x axis) contains every observed proportion for that cell type across all samples in donor 15496’s brain. Right column are the spatial correlations between cell type proportions between datasets (p-values in Supplementary Table 2).

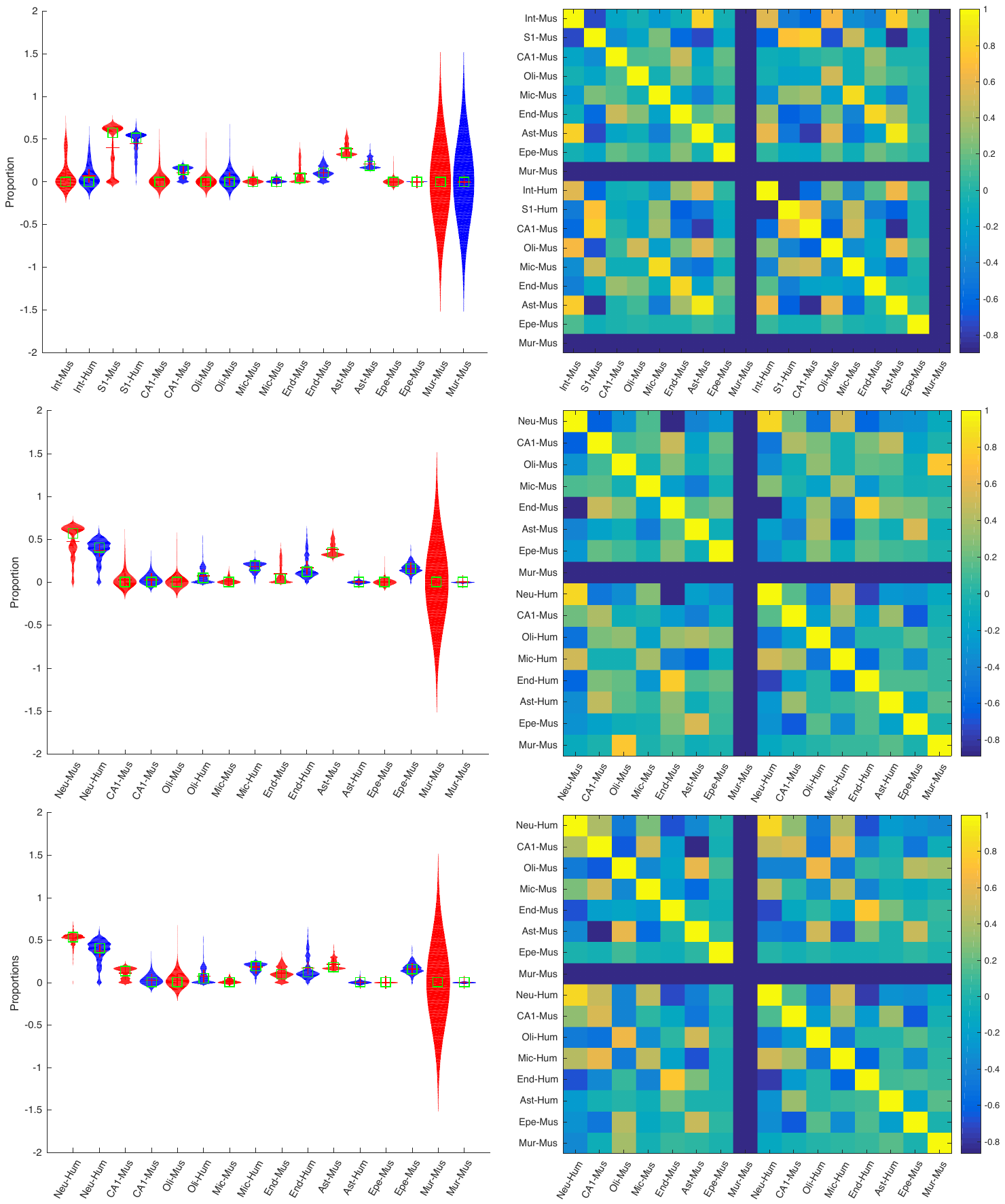

Supplementary Figure S10 Comparison of cell type distributions between each combination of datasets within donor 15697. Top: MusNG (red) and HumN (blue), Middle: MusNG (red) and HumNG (blue), Bottom: HumN (red) and HumNG (blue). Left column are the cell type distributions from donor 15697 such that each cell type signature distribution (x axis) contains every observed proportion for that cell type across all samples in donor 15697’s brain. Right column are the spatial correlations between cell type proportions between datasets (p-values in Supplementary Table 2).

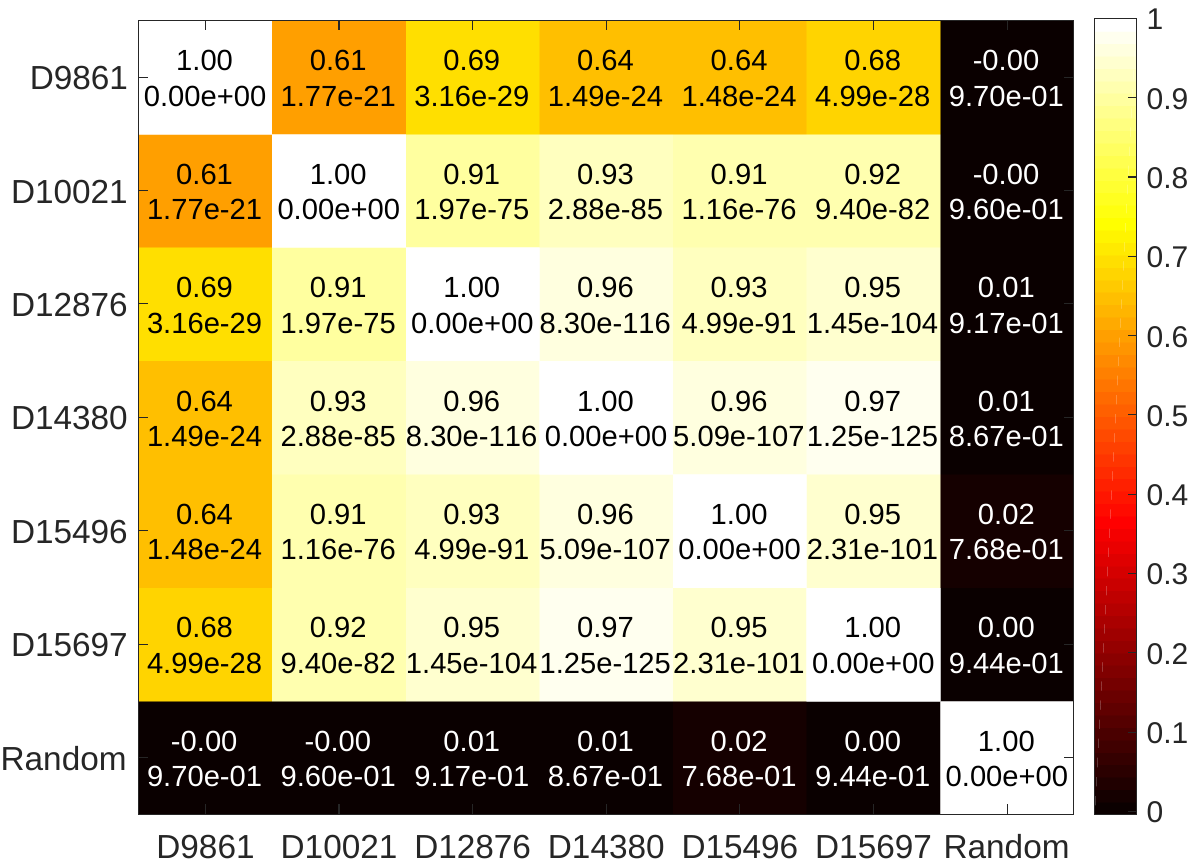

Supplementary Figure S11 Donor to donor comparison using MusNG expression signatures. The cell type distributions across all 6 brains were compared by pooling the samples into overlapping anatomic locations. The random dataset constitutes a five-fold randomization of the sample labels such that it reflects the background noise and spurious correlation. Each row and column are a brain donor. Anova p-value between random and non-random donors is 1.1799×10­^-23^.

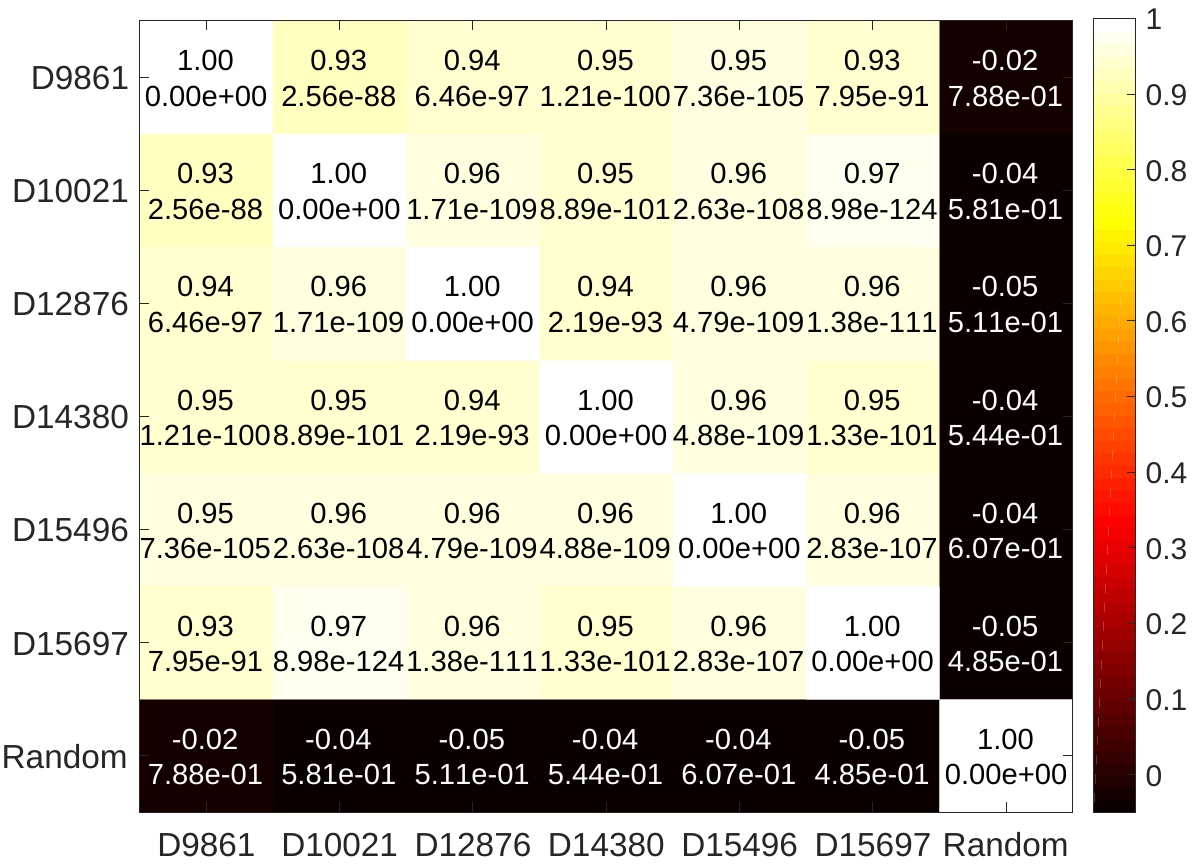

Supplementary Figure S12 Donor to donor comparison using HumN expression signatures. The cell type distributions across all 6 brains were compared by pooling the samples into overlapping anatomic locations. The random dataset constitutes a five-fold randomization of the sample labels such that it reflects the background noise and spurious correlation. Each row and column are a brain donor. Anova p-value between random and non-random donors is 1.5364×10­^-27^.

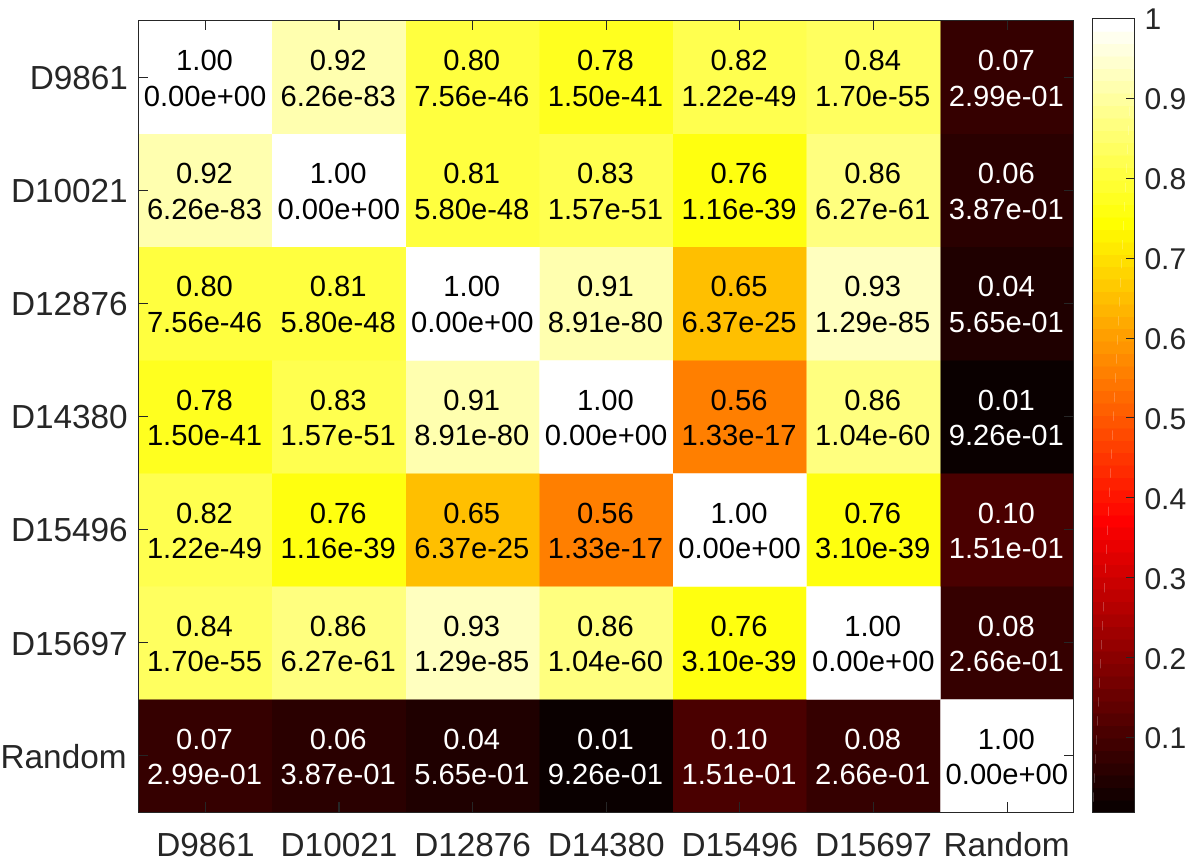

Supplementary Figure S13 Donor to donor comparison using HumNG expression signatures. The cell type distributions across all 6 brains were compared by pooling the samples into overlapping anatomic locations. The random dataset constitutes a five-fold randomization of the sample labels such that it reflects the background noise and spurious correlation. Each row and column are a brain donor. Anova p-value between random and non-random donors is 2.1421×10­^-11^.

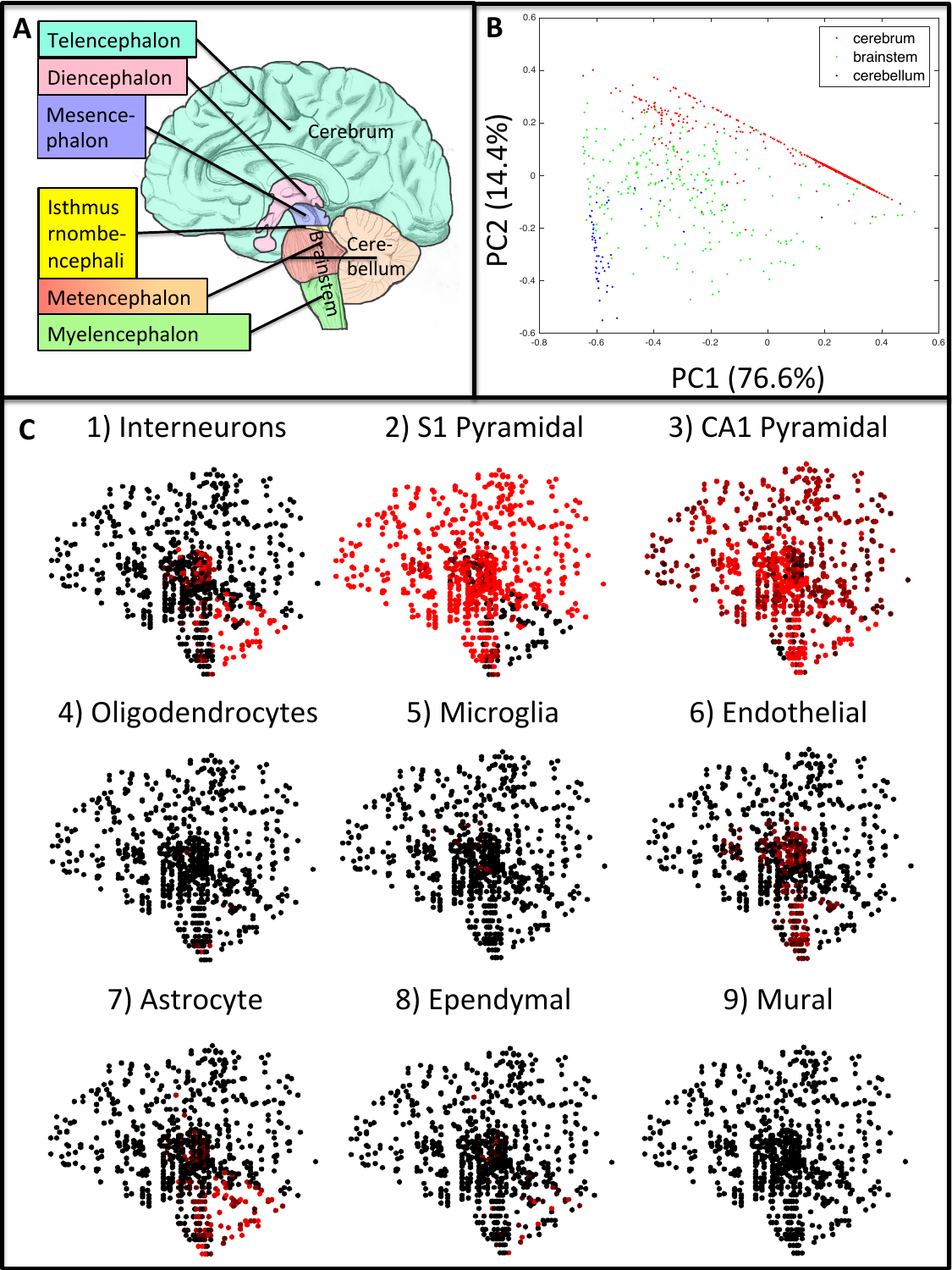

Supplementary Figure S14 Example of spatial distributions of cell type proportions in AHBA donor 9861 using MusNG to deconvolute the RNA expression profile. A) A reference map with developmental structures and major brain regions marked. B) PCA plot of the cell type proportion matrix such that the 9 cell types are reduced to 2 principal cell types for each of the samples in the brain. The colors indicate the three regions (cerebrum, brainstem, and cerebellum). C) The proportion of each cell type plotted from a sagittal view; Red: high cell proportion (1); Black: low cell proportion (0) on a scale 0-1.

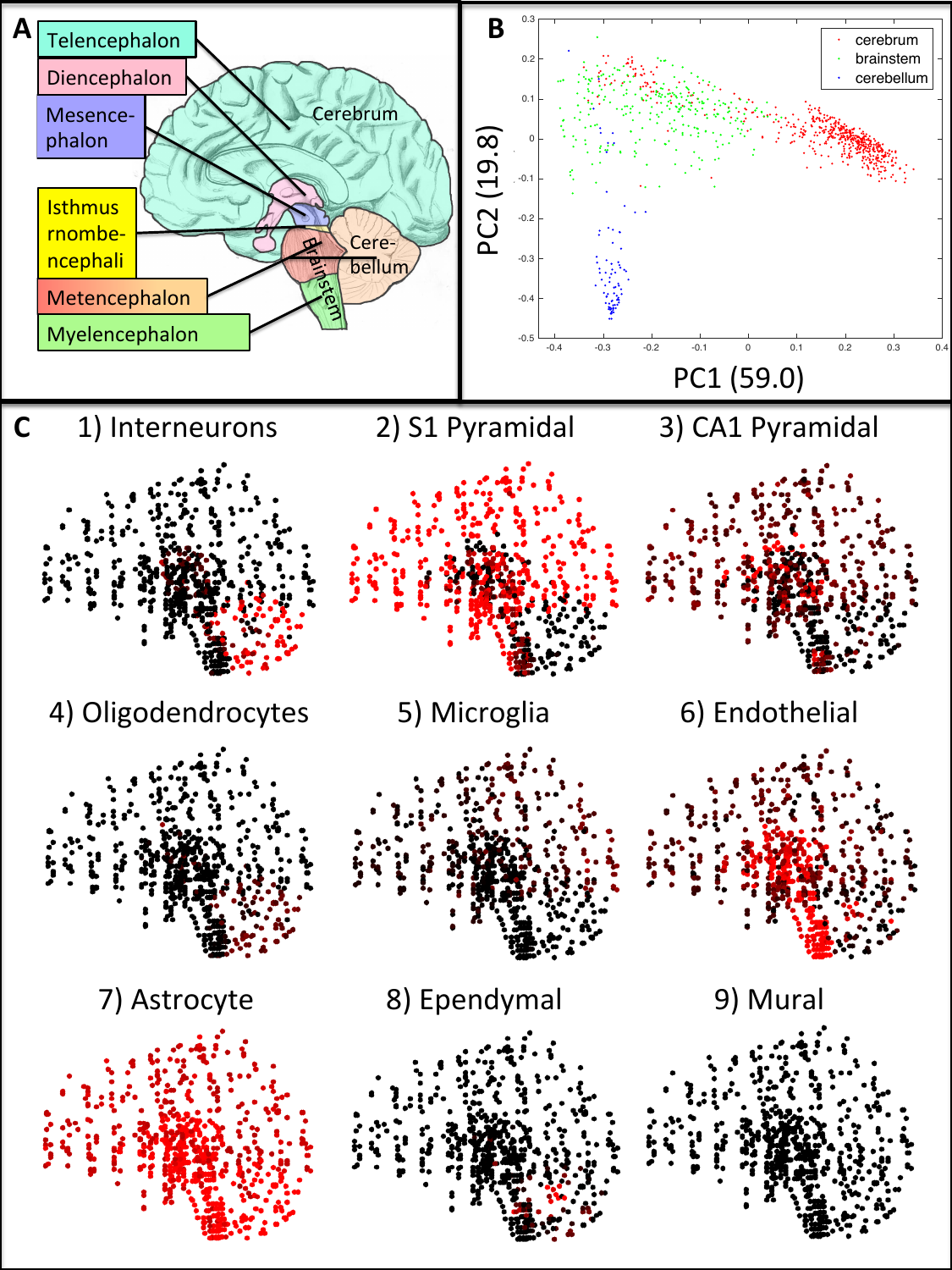

Supplementary Figure S15 Example of spatial distributions of cell type proportions in AHBA donor 10021 using MusNG to deconvolute the RNA expression profile. A) A reference map with developmental structures and major brain regions marked. B) PCA plot of the cell type proportion matrix such that the 9 cell types are reduced to 2 principal cell types for each of the samples in the brain. The colors indicate the three regions (cerebrum, brainstem, and cerebellum). C) The proportion of each cell type plotted from a sagittal view; Red: high cell proportion (1); Black: low cell proportion (0) on a scale 0-1.

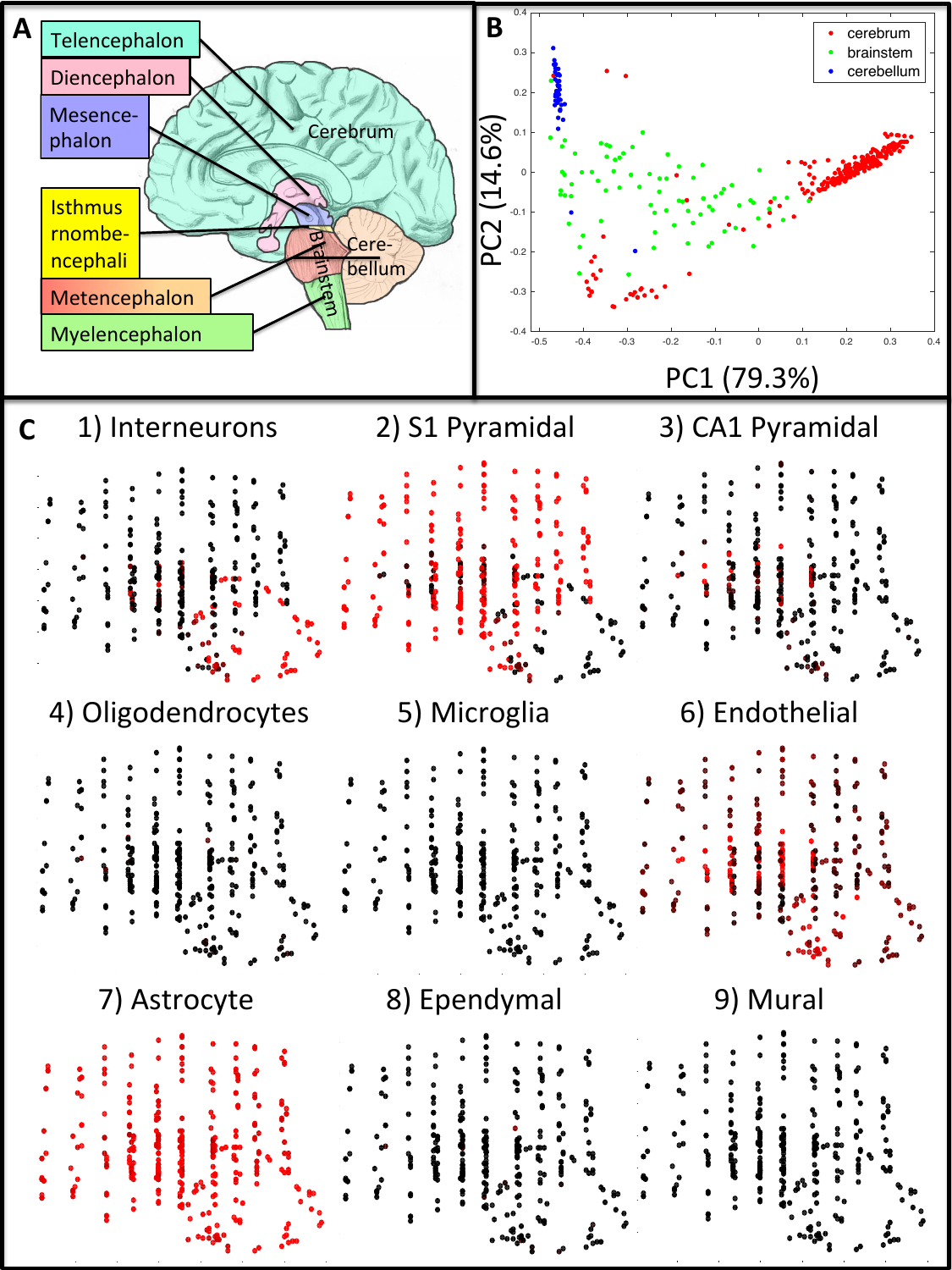

Supplementary Figure S16 Example of spatial distributions of cell type proportions in AHBA donor 12876 using MusNG to deconvolute the RNA expression profile. A) A reference map with developmental structures and major brain regions marked. B) PCA plot of the cell type proportion matrix such that the 9 cell types are reduced to 2 principal cell types for each of the samples in the brain. The colors indicate the three regions (cerebrum, brainstem, and cerebellum). C) The proportion of each cell type plotted from a sagittal view; Red: high cell proportion (1); Black: low cell proportion (0) on a scale 0-1.

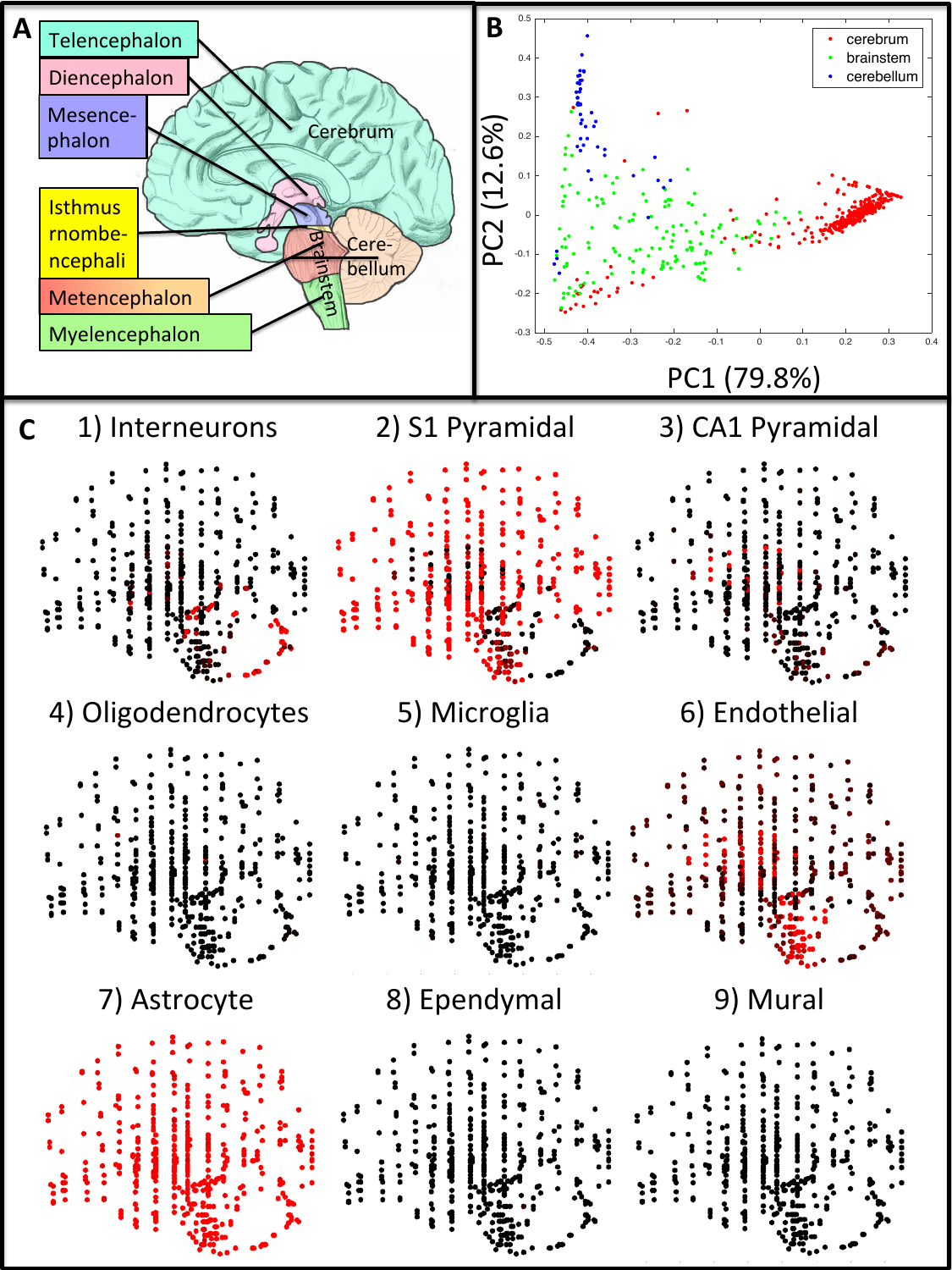

Supplementary Figure S17 Example of spatial distributions of cell type proportions in AHBA donor 14380 using MusNG to deconvolute the RNA expression profile. A) A reference map with developmental structures and major brain regions marked. B) PCA plot of the cell type proportion matrix such that the 9 cell types are reduced to 2 principal cell types for each of the samples in the brain. The colors indicate the three regions (cerebrum, brainstem, and cerebellum). C) The proportion of each cell type plotted from a sagittal view; Red: high cell proportion (1); Black: low cell proportion (0) on a scale 0-1.

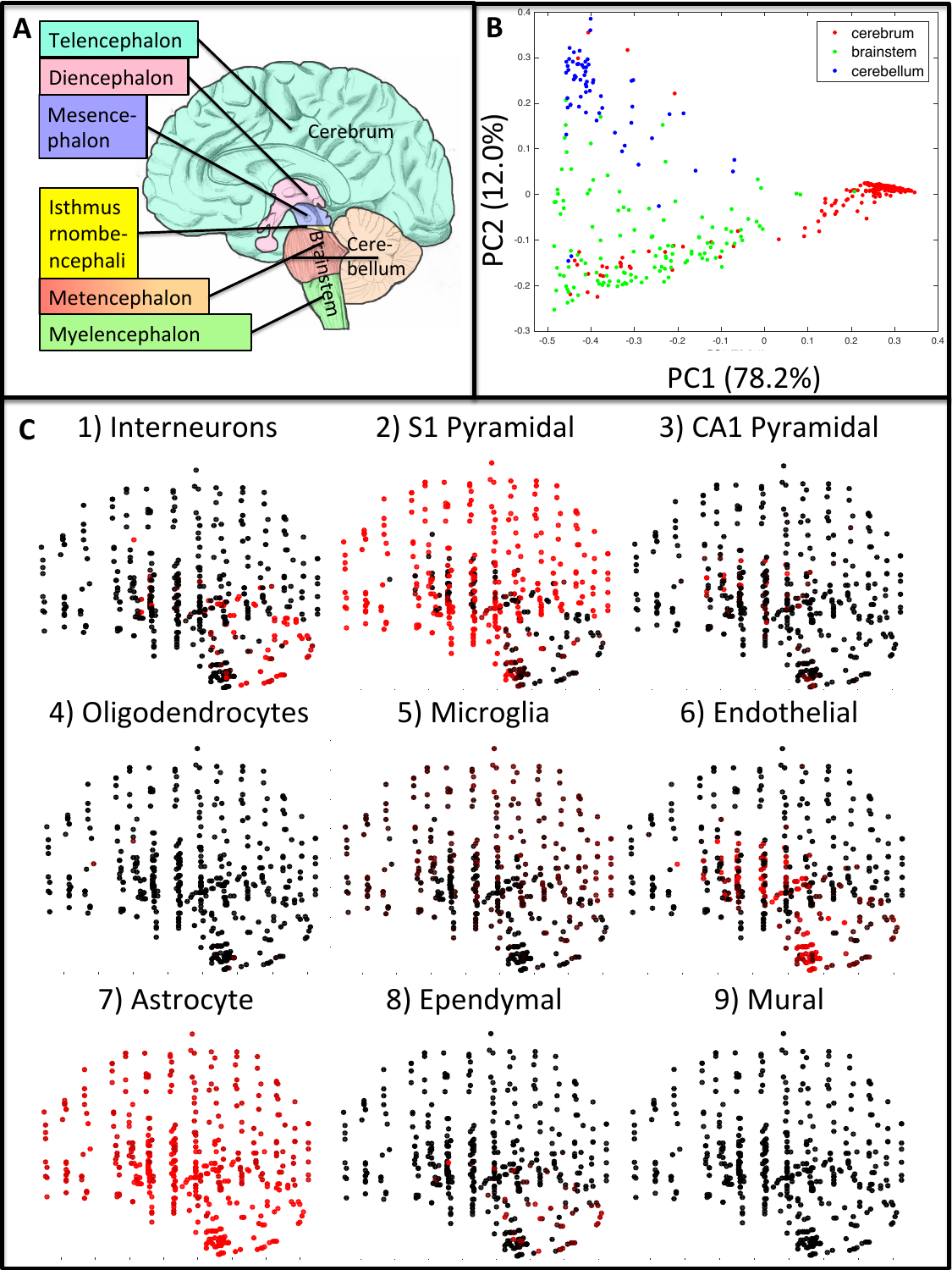

Supplementary Figure S18 Example of spatial distributions of cell type proportions in AHBA donor 15496 using MusNG to deconvolute the RNA expression profile. A) A reference map with developmental structures and major brain regions marked. B) PCA plot of the cell type proportion matrix such that the 9 cell types are reduced to 2 principal cell types for each of the samples in the brain. The colors indicate the three regions (cerebrum, brainstem, and cerebellum). C) The proportion of each cell type plotted from a sagittal view; Red: high cell proportion (1); Black: low cell proportion (0) on a scale 0-1.

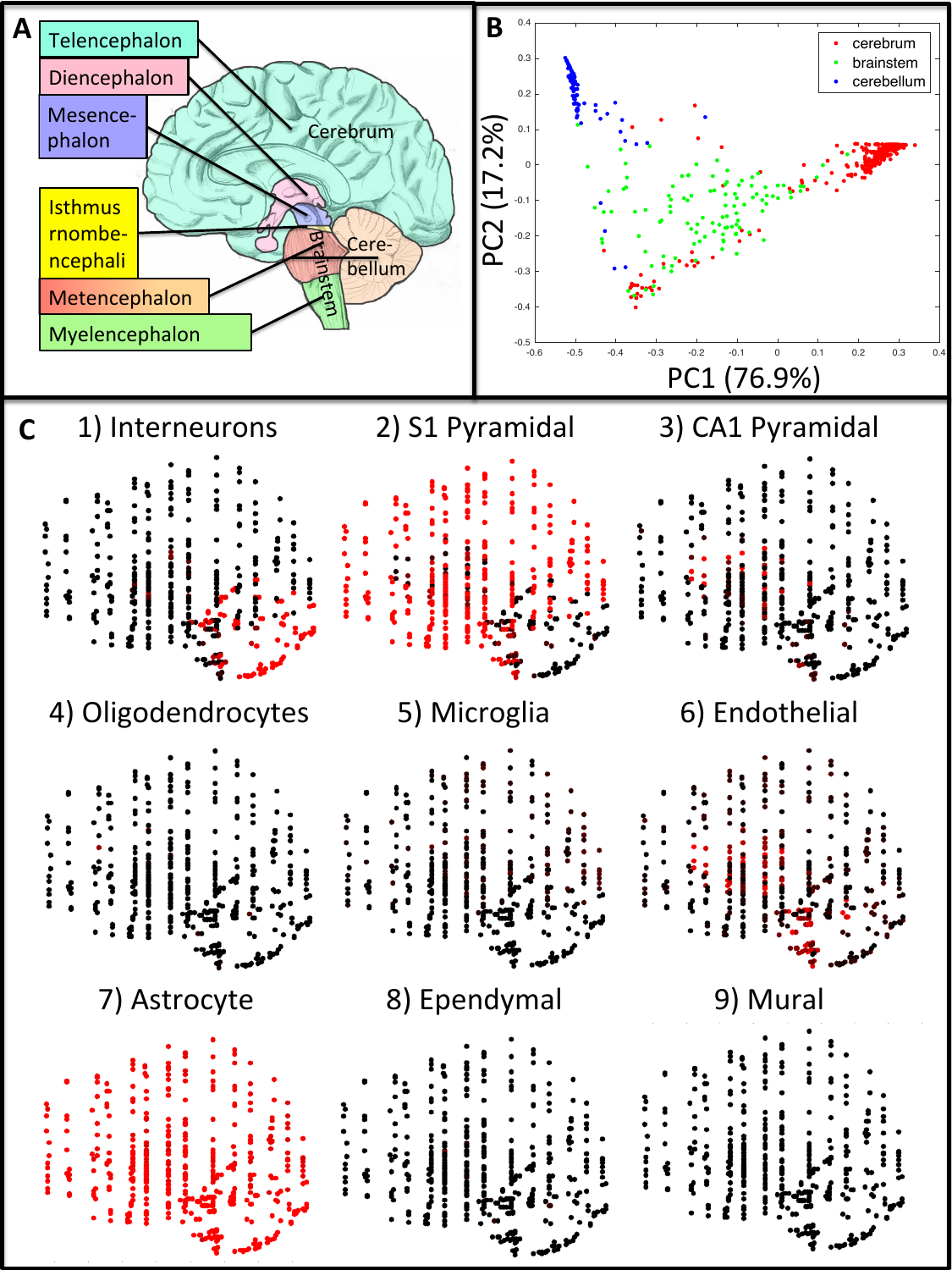

Supplementary Figure S19 Example of spatial distributions of cell type proportions in AHBA donor 15697 using MusNG to deconvolute the RNA expression profile. A) A reference map with developmental structures and major brain regions marked. B) PCA plot of the cell type proportion matrix such that the 9 cell types are reduced to 2 principal cell types for each of the samples in the brain. The colors indicate the three regions (cerebrum, brainstem, and cerebellum). C) The proportion of each cell type plotted from a sagittal view; Red: high cell proportion (1); Black: low cell proportion (0) on a scale 0-1.

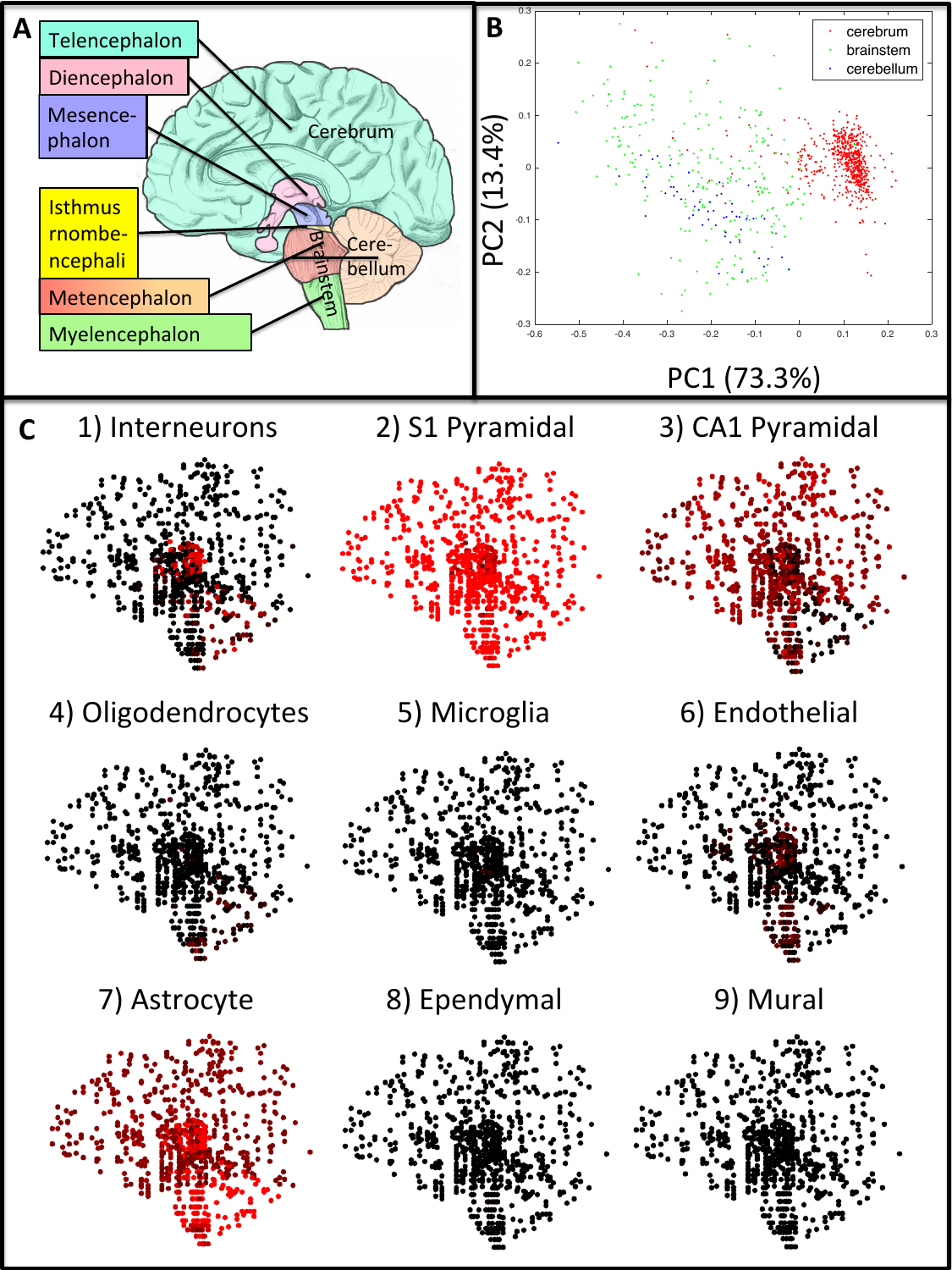

Supplementary Figure S20 Example of spatial distributions of cell type proportions in AHBA donor 9861 using HumN to deconvolute the RNA expression profile. A) A reference map with developmental structures and major brain regions marked. B) PCA plot of the cell type proportion matrix such that the 9 cell types are reduced to 2 principal cell types for each of the samples in the brain. The colors indicate the three regions (cerebrum, brainstem, and cerebellum). C) The proportion of each cell type plotted from a sagittal view; Red: high cell proportion (1); Black: low cell proportion (0) on a scale 0-1.

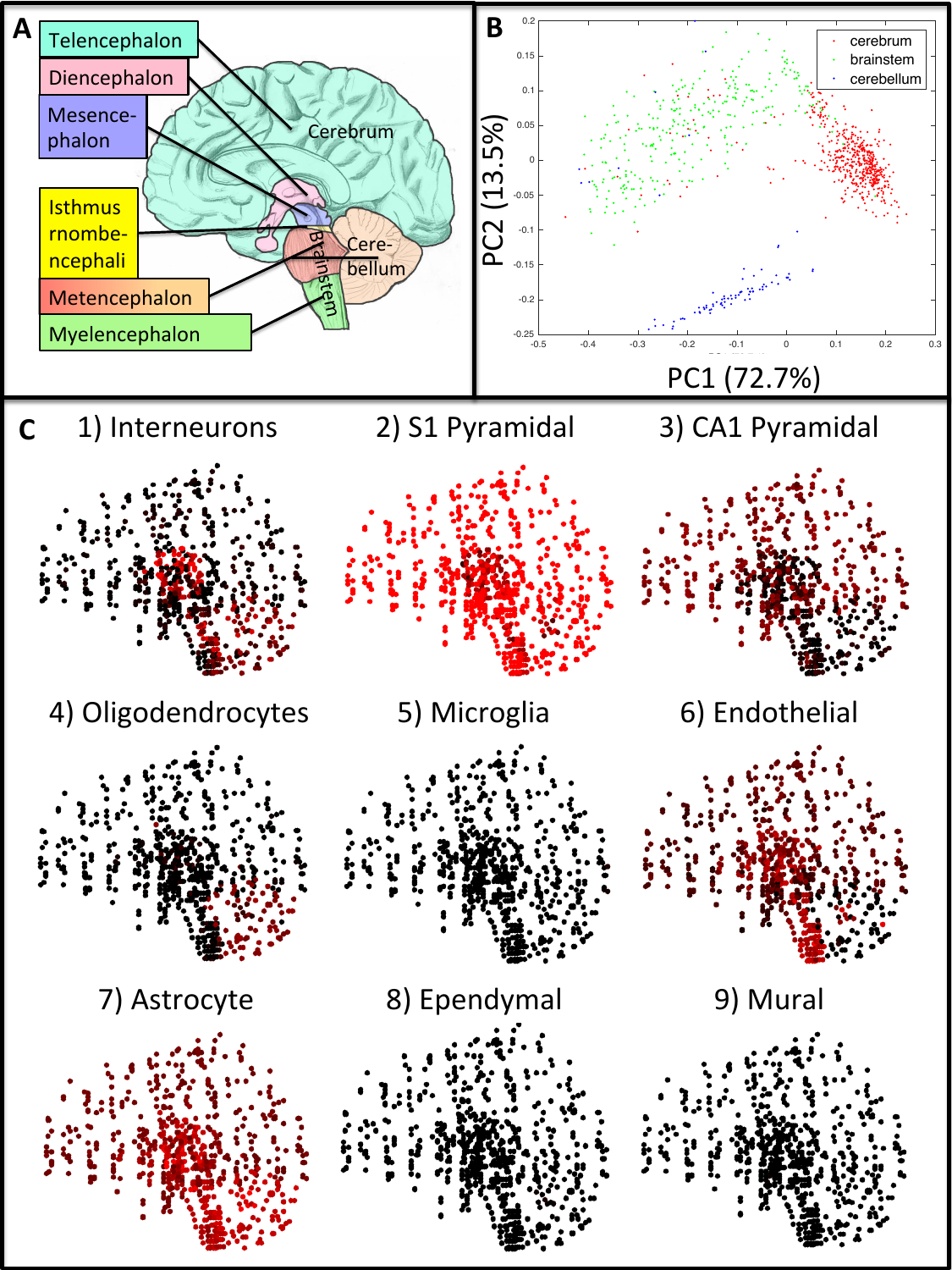

Supplementary Figure S21 Example of spatial distributions of cell type proportions in AHBA donor 10021 using HumN to deconvolute the RNA expression profile. A) A reference map with developmental structures and major brain regions marked. B) PCA plot of the cell type proportion matrix such that the 9 cell types are reduced to 2 principal cell types for each of the samples in the brain. The colors indicate the three regions (cerebrum, brainstem, and cerebellum). C) The proportion of each cell type plotted from a sagittal view; Red: high cell proportion (1); Black: low cell proportion (0) on a scale 0-1.

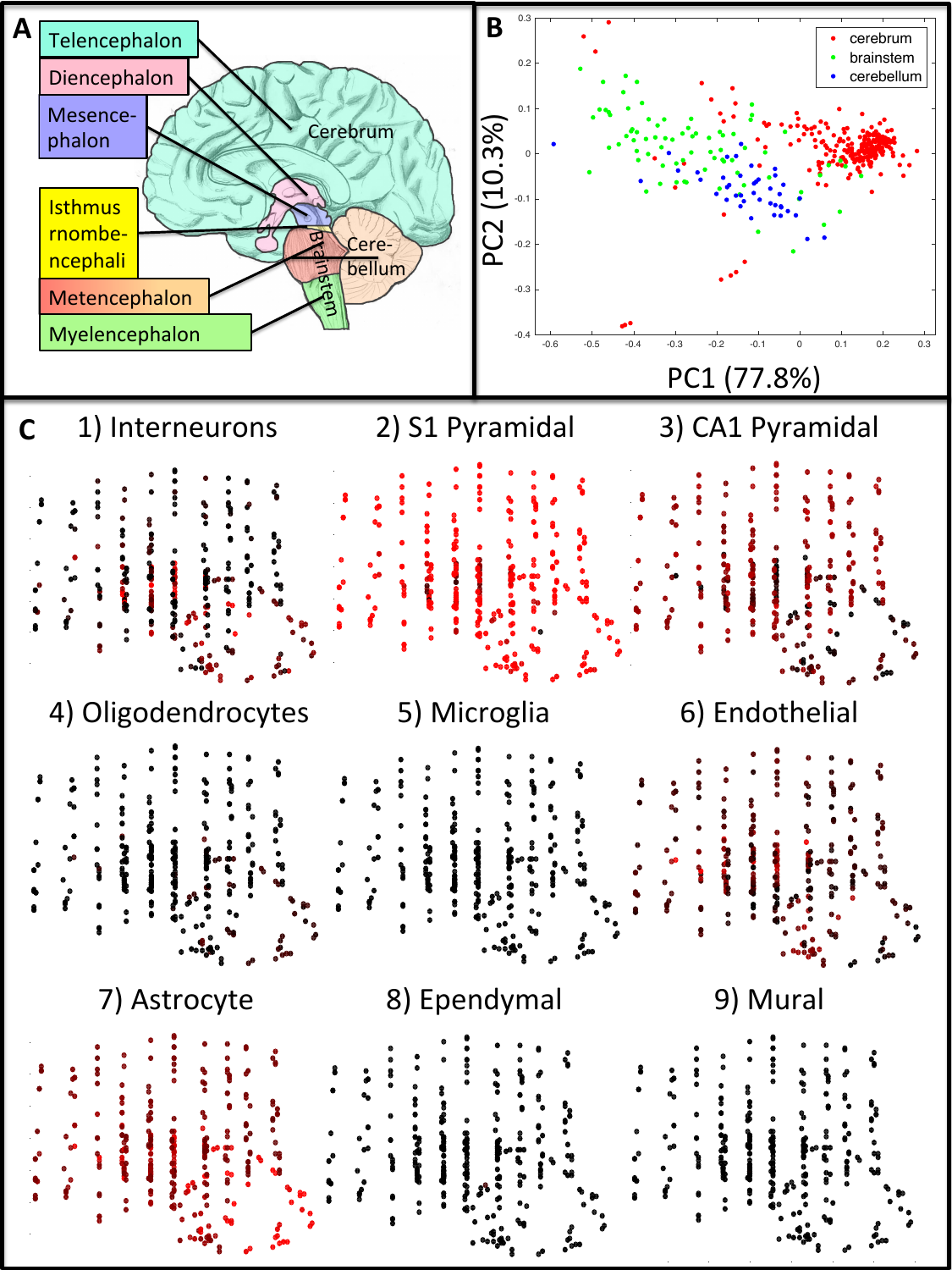

Supplementary Figure S22 Example of spatial distributions of cell type proportions in AHBA donor 12876 using HumN to deconvolute the RNA expression profile. A) A reference map with developmental structures and major brain regions marked. B) PCA plot of the cell type proportion matrix such that the 9 cell types are reduced to 2 principal cell types for each of the samples in the brain. The colors indicate the three regions (cerebrum, brainstem, and cerebellum). C) The proportion of each cell type plotted from a sagittal view; Red: high cell proportion (1); Black: low cell proportion (0) on a scale 0-1.

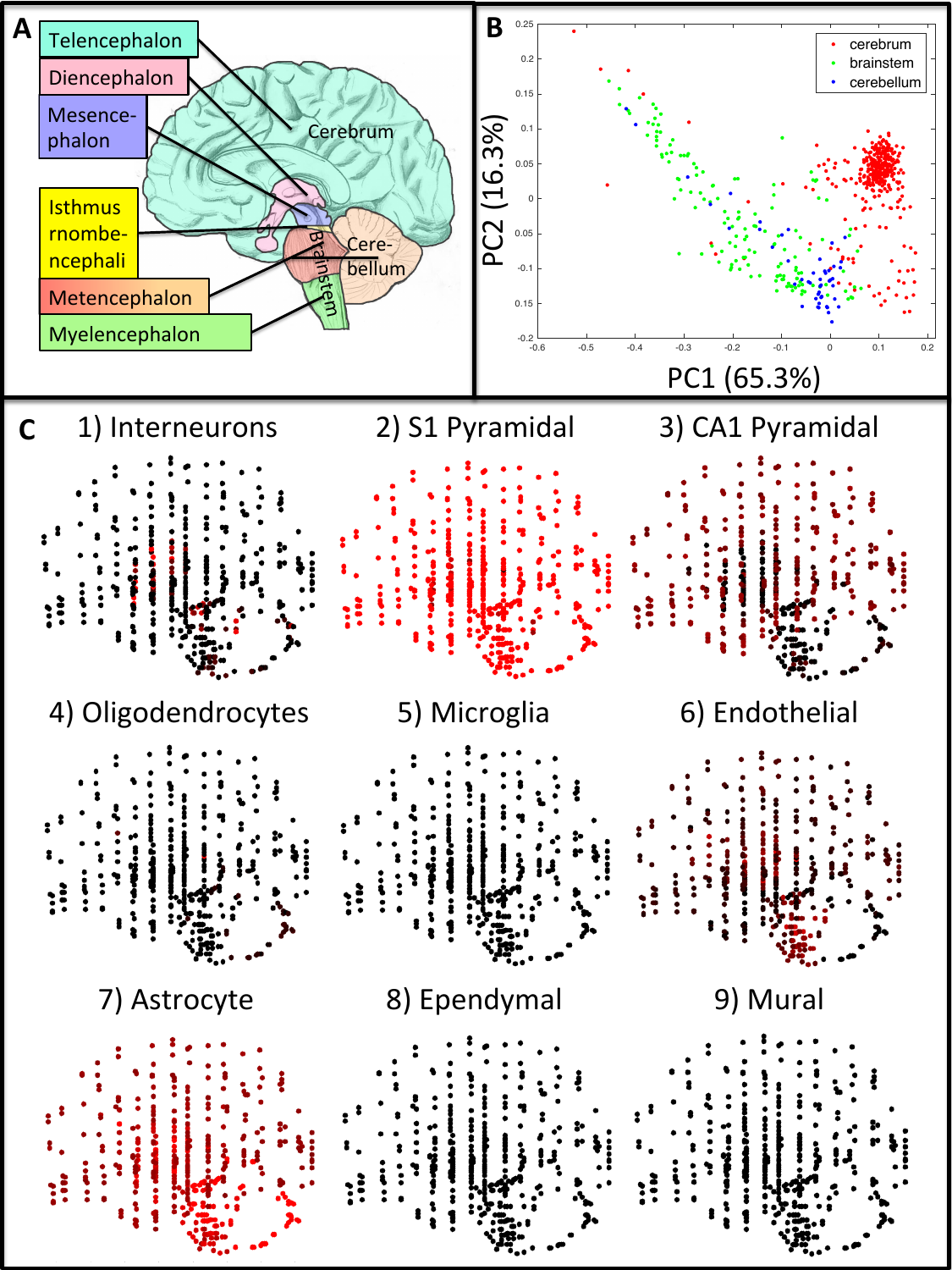

Supplementary Figure S23 Example of spatial distributions of cell type proportions in AHBA donor 14380 using HumN to deconvolute the RNA expression profile. A) A reference map with developmental structures and major brain regions marked. B) PCA plot of the cell type proportion matrix such that the 9 cell types are reduced to 2 principal cell types for each of the samples in the brain. The colors indicate the three regions (cerebrum, brainstem, and cerebellum). C) The proportion of each cell type plotted from a sagittal view; Red: high cell proportion (1); Black: low cell proportion (0) on a scale 0-1.

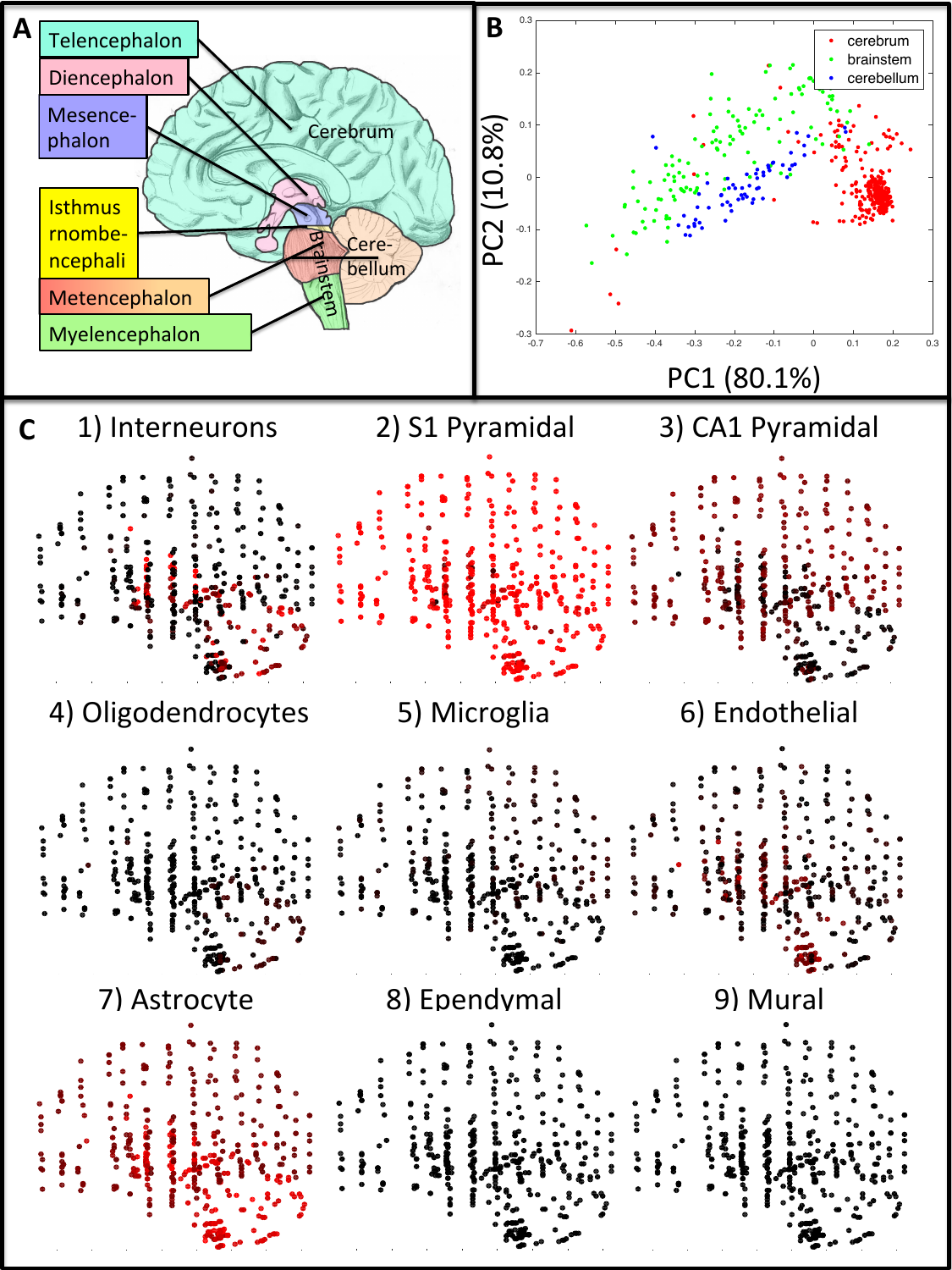

Supplementary Figure S24 Example of spatial distributions of cell type proportions in AHBA donor 15496 using HumN to deconvolute the RNA expression profile. A) A reference map with developmental structures and major brain regions marked. B) PCA plot of the cell type proportion matrix such that the 9 cell types are reduced to 2 principal cell types for each of the samples in the brain. The colors indicate the three regions (cerebrum, brainstem, and cerebellum). C) The proportion of each cell type plotted from a sagittal view; Red: high cell proportion (1); Black: low cell proportion (0) on a scale 0-1.

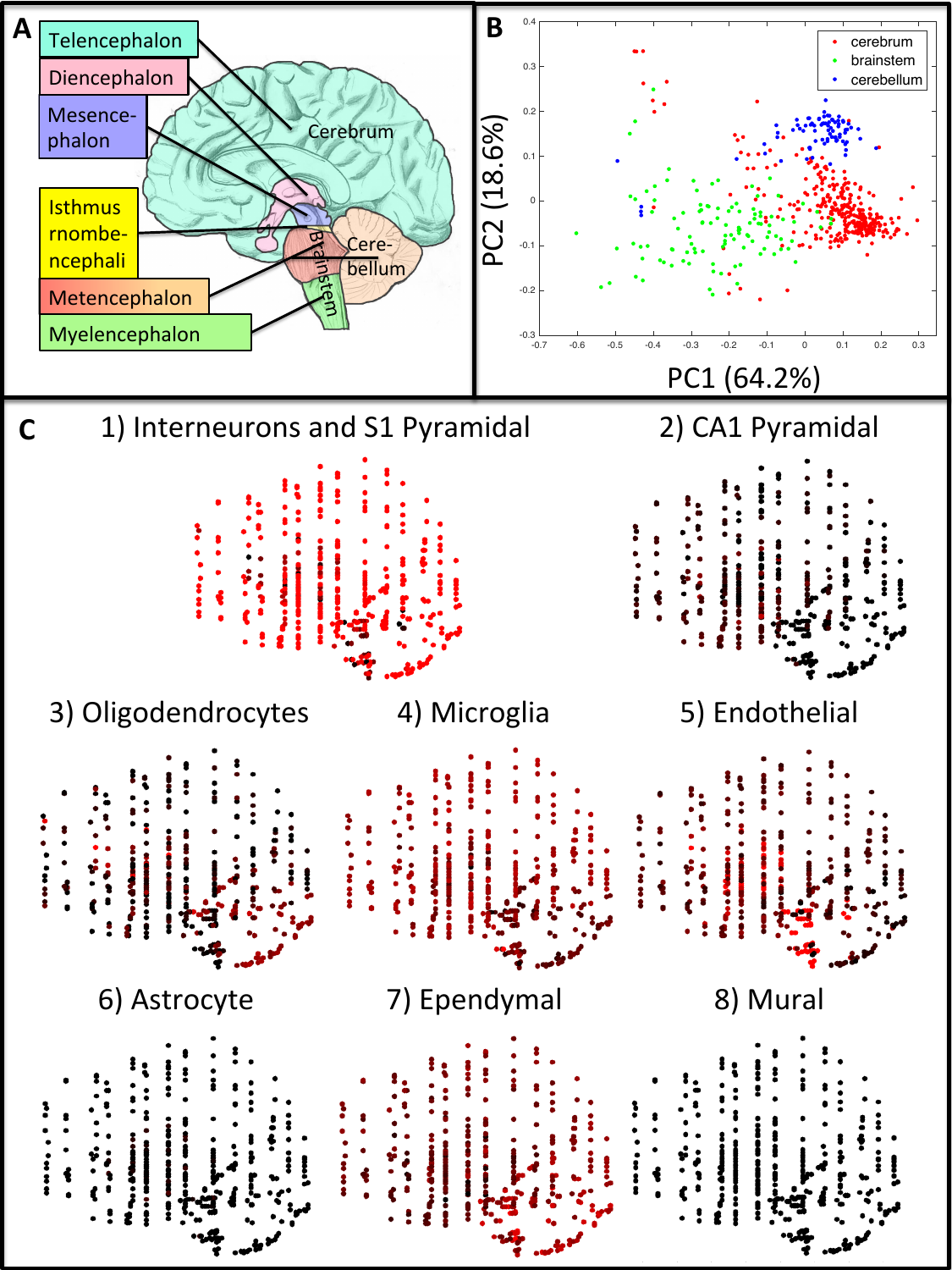

Supplementary Figure S25 Example of spatial distributions of cell type proportions in AHBA donor 15697 using HumN to deconvolute the RNA expression profile. A) A reference map with developmental structures and major brain regions marked. B) PCA plot of the cell type proportion matrix such that the 9 cell types are reduced to 2 principal cell types for each of the samples in the brain. The colors indicate the three regions (cerebrum, brainstem, and cerebellum). C) The proportion of each cell type plotted from a sagittal view; Red: high cell proportion (1); Black: low cell proportion (0) on a scale 0-1.

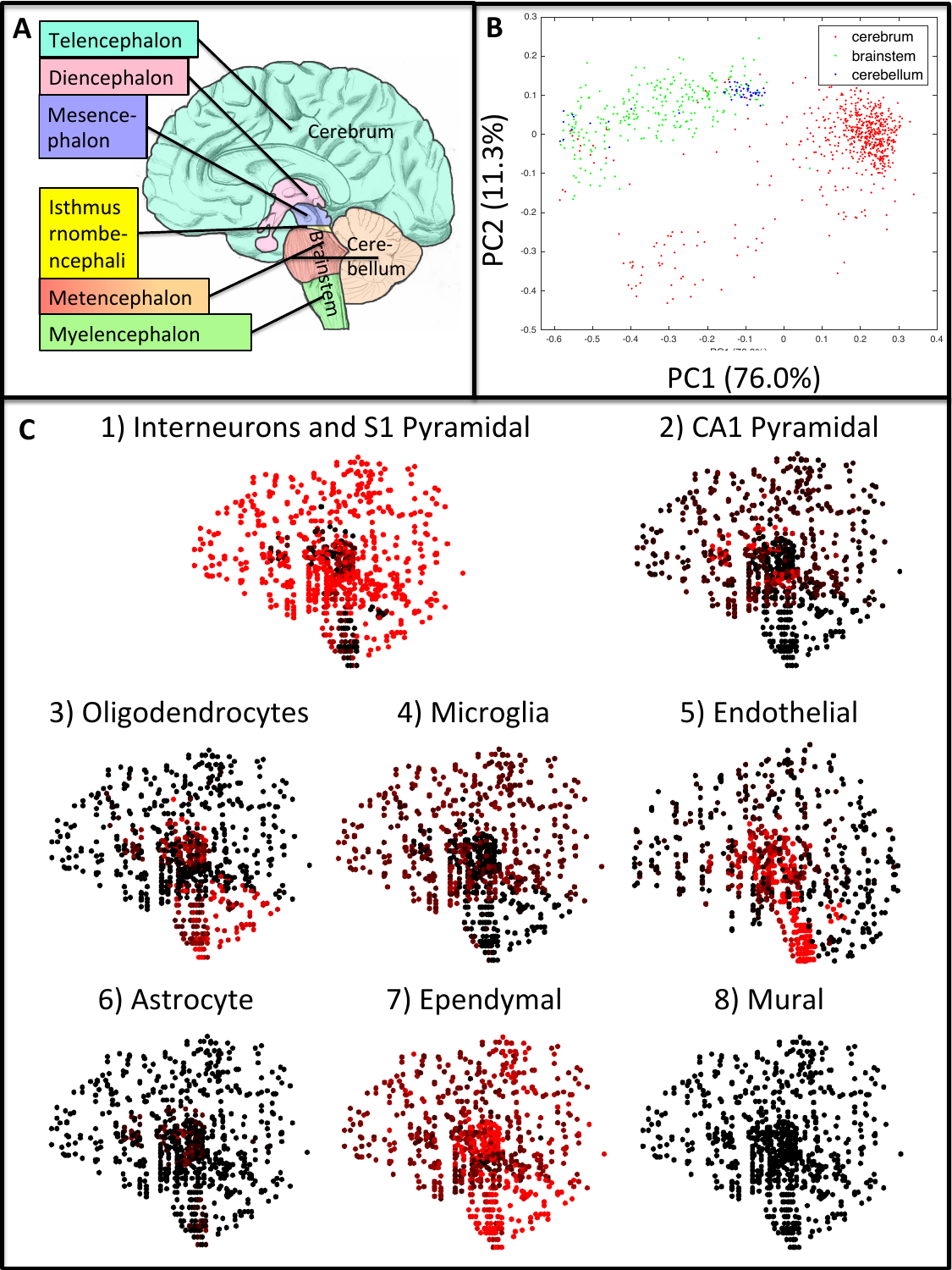

Supplementary Figure S26 Example of spatial distributions of cell type proportions in AHBA donor 9861 using HumNG to deconvolute the RNA expression profile. A) A reference map with developmental structures and major brain regions marked. B) PCA plot of the cell type proportion matrix such that the 9 cell types are reduced to 2 principal cell types for each of the samples in the brain. The colors indicate the three regions (cerebrum, brainstem, and cerebellum). C) The proportion of each cell type plotted from a sagittal view; Red: high cell proportion (1); Black: low cell proportion (0) on a scale 0-1.

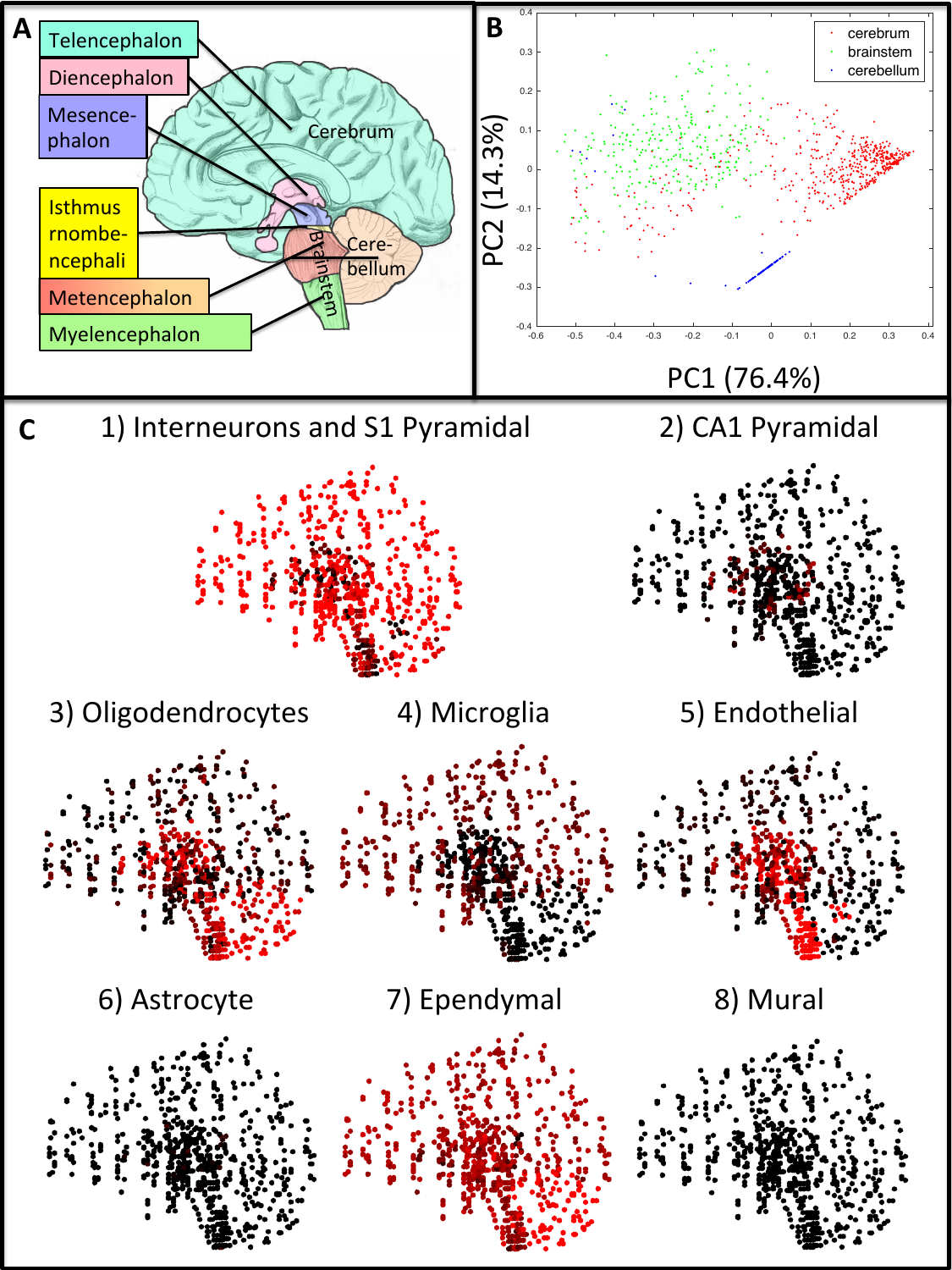

Supplementary Figure S27 Example of spatial distributions of cell type proportions in AHBA donor 10021 using HumNG to deconvolute the RNA expression profile. A) A reference map with developmental structures and major brain regions marked. B) PCA plot of the cell type proportion matrix such that the 9 cell types are reduced to 2 principal cell types for each of the samples in the brain. The colors indicate the three regions (cerebrum, brainstem, and cerebellum). C) The proportion of each cell type plotted from a sagittal view; Red: high cell proportion (1); Black: low cell proportion (0) on a scale 0-1.

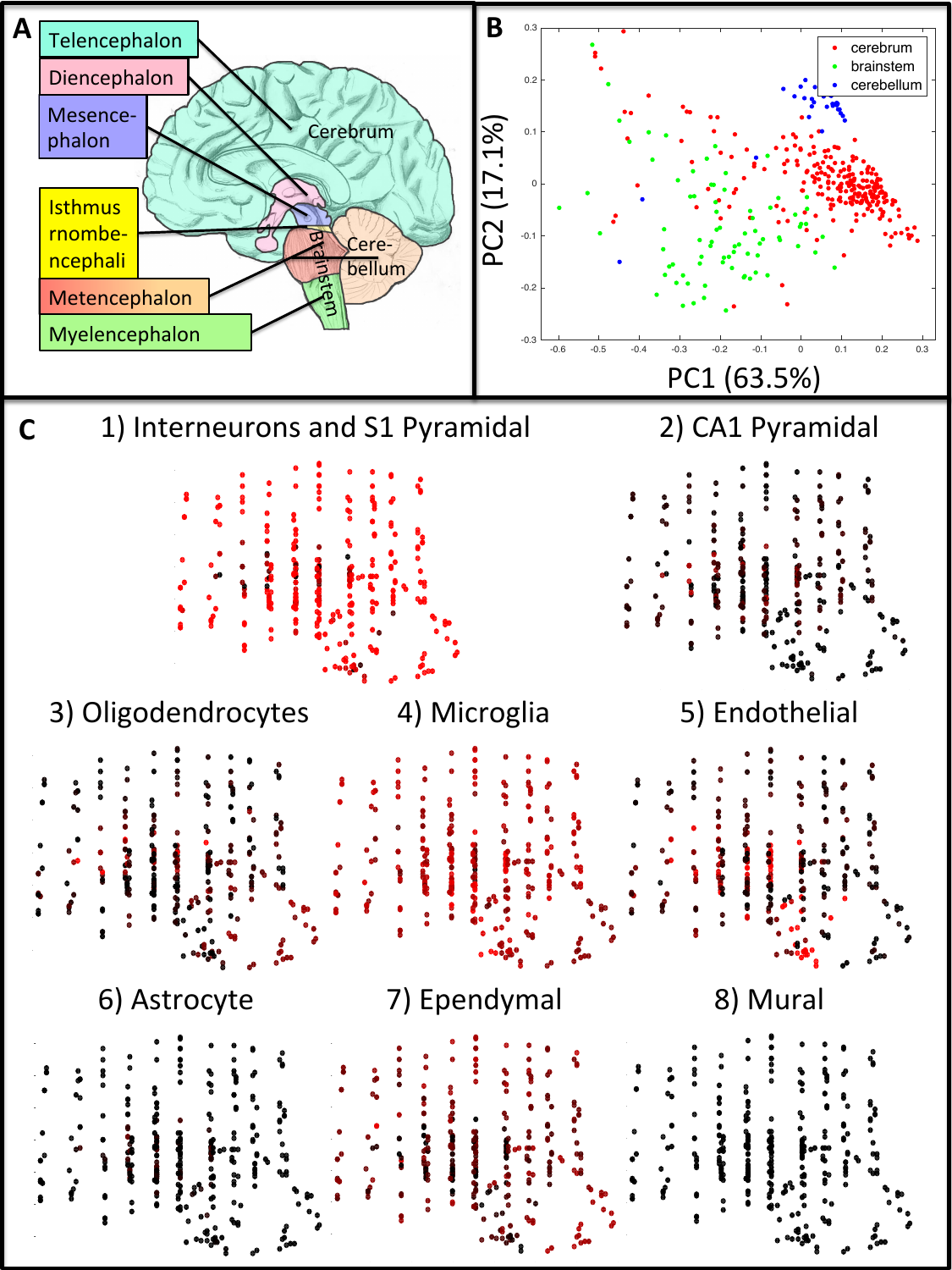

Supplementary Figure S28 Example of spatial distributions of cell type proportions in AHBA donor 12876 using HumNG to deconvolute the RNA expression profile. A) A reference map with developmental structures and major brain regions marked. B) PCA plot of the cell type proportion matrix such that the 9 cell types are reduced to 2 principal cell types for each of the samples in the brain. The colors indicate the three regions (cerebrum, brainstem, and cerebellum). C) The proportion of each cell type plotted from a sagittal view; Red: high cell proportion (1); Black: low cell proportion (0) on a scale 0-1.

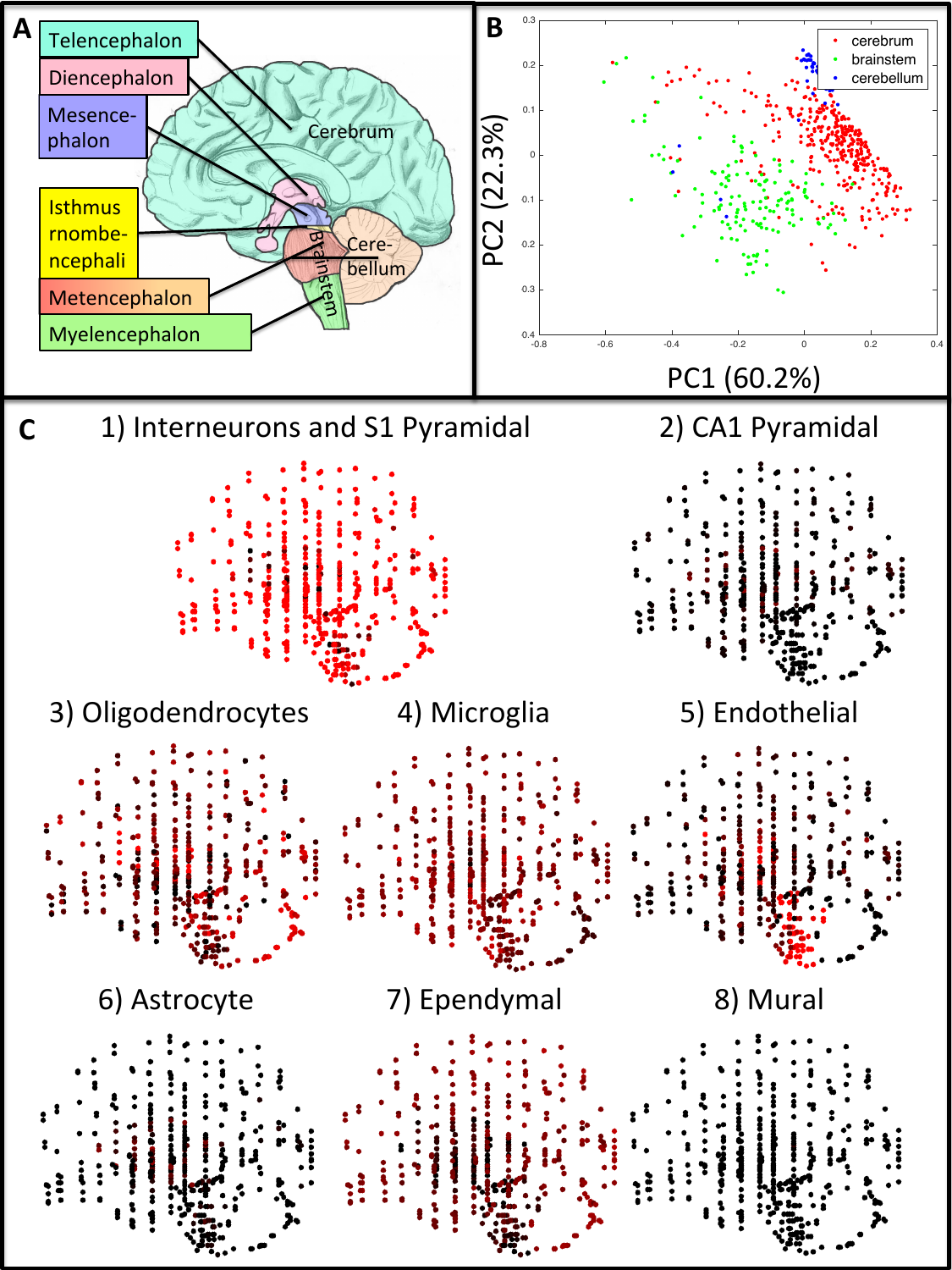

Supplementary Figure S29 Example of spatial distributions of cell type proportions in AHBA donor 14380 using HumNG to deconvolute the RNA expression profile. A) A reference map with developmental structures and major brain regions marked. B) PCA plot of the cell type proportion matrix such that the 9 cell types are reduced to 2 principal cell types for each of the samples in the brain. The colors indicate the three regions (cerebrum, brainstem, and cerebellum). C) The proportion of each cell type plotted from a sagittal view; Red: high cell proportion (1); Black: low cell proportion (0) on a scale 0-1.

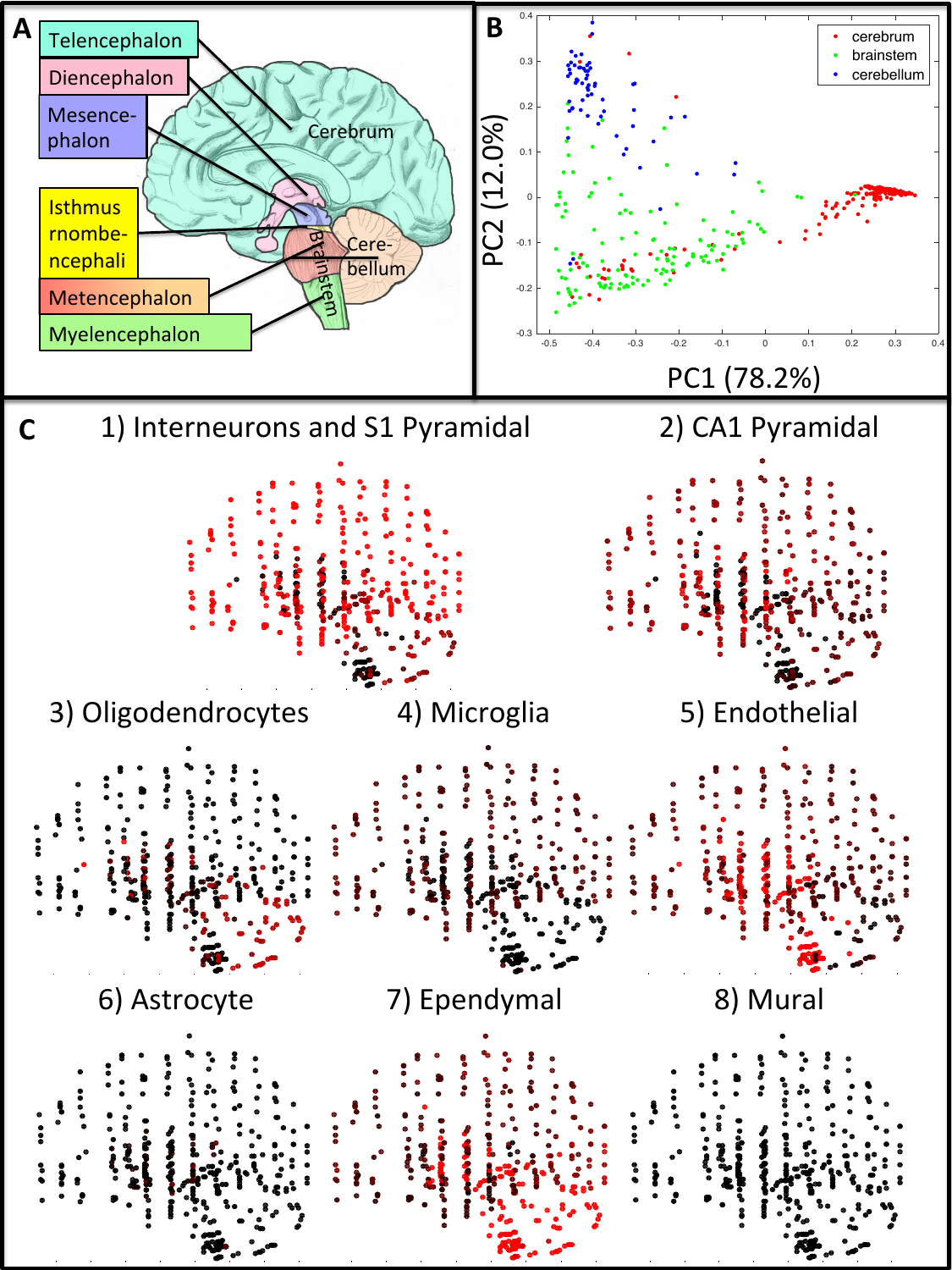

Supplementary Figure S30 Example of spatial distributions of cell type proportions in AHBA donor 15496 using HumNG to deconvolute the RNA expression profile. A) A reference map with developmental structures and major brain regions marked. B) PCA plot of the cell type proportion matrix such that the 9 cell types are reduced to 2 principal cell types for each of the samples in the brain. The colors indicate the three regions (cerebrum, brainstem, and cerebellum). C) The proportion of each cell type plotted from a sagittal view; Red: high cell proportion (1); Black: low cell proportion (0) on a scale 0-1.

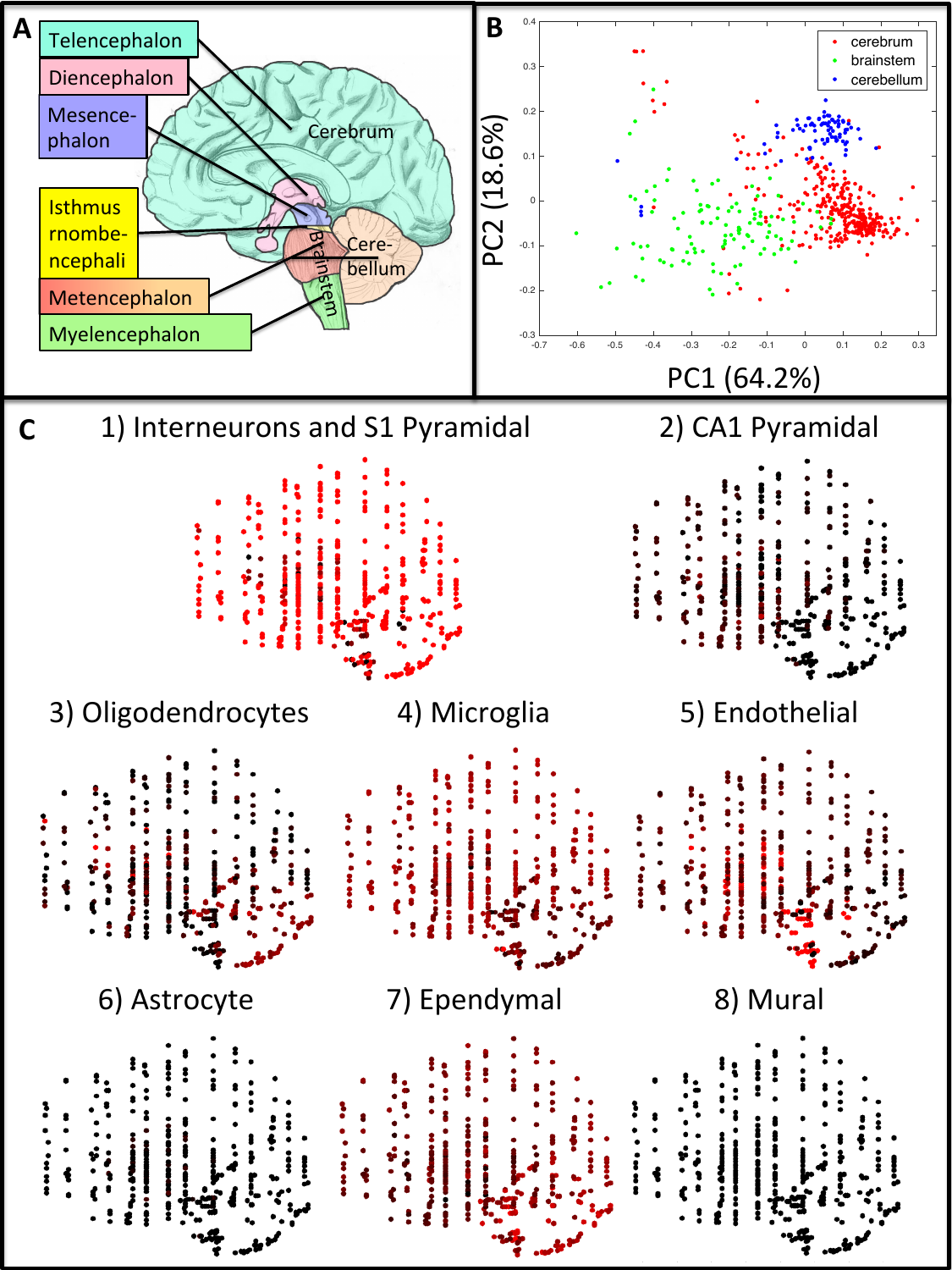

Supplementary Figure S31 Example of spatial distributions of cell type proportions in AHBA donor 15697 using HumNG to deconvolute the RNA expression profile. A) A reference map with developmental structures and major brain regions marked. B) PCA plot of the cell type proportion matrix such that the 9 cell types are reduced to 2 principal cell types for each of the samples in the brain. The colors indicate the three regions (cerebrum, brainstem, and cerebellum). C) The proportion of each cell type plotted from a sagittal view; Red: high cell proportion (1); Black: low cell proportion (0) on a scale 0-1.

|  | D9861 | D10021 | D12876 | D14380 | D15496 | D15697 |
| --- | --- | --- | --- | --- | --- | --- |
| MusNG-HumN | 8.5415×10^-15^ | 2.1110×10^-14^ | 9.4471×10^-12^ | 9.2133×10^-12^ | 3.2302×10^-08^ | 1.5885×10^-09^ |
| MusNG-HumNG | 2.5544×10^-04^ | 5.1148×10^-10^ | 8.7296×10^-06^ | 5.9451×10^-06^ | 1.5071×10^-05^ | 1.9738×10^-06^ |
| HumN-HumNG | 3.0739×10^-05^ | 7.8195×10^-06^ | 7.1236×10^-08^ | 6.6163×10^-06^ | 3.3409×10^-06^ | 1.2331×10^-08^ |

Supplementary Table S1 Each row is the combination scRNA-Seq datasets being compared. Each column is a brain donor. The cells contain p-values from an anova of cell type correlation values grouped by 1) same cell type-between datasets versus 2) different cell type-between datasets (within dataset same cell type correlations removed because they have a correlation of 1). See Consistency of Cell Type Proportions Between Datasets in the results section for a detailed description of how these p-values were obtained.

|  | Accuracy - overall | Sens-cerebrum | Sens - brainstem | Sens - cerebellum | Spec – cerebrum | Spec – brainstem | Spec - cerebellumq | Fscore –cerebrum | Fscore – brainstem | Fscore - cerebellum |
| --- | --- | --- | --- | --- | --- | --- | --- | --- | --- | --- |
| MusNG-d9861 | 0.7030 | 0.7994 | 0.4080 | 0.9245 | 0.8581 | 0.8132 | 0.8791 | 0.8567 | 0.4232 | 0.4667 |
| MusNG-d10021 | 0.8813 | 0.8409 | 0.9539 | 0.8916 | 0.9671 | 0.8478 | 0.9988 | 0.9024 | 0.8354 | 0.9367 |
| MusNG-d12876 | 0.8182 | 0.8642 | 0.6026 | 0.9524 | 0.9500 | 0.8877 | 0.9128 | 0.9150 | 0.5987 | 0.7273 |
| MusNG-d14380 | 0.8696 | 0.9021 | 0.8056 | 0.8333 | 0.9479 | 0.9039 | 0.9543 | 0.9339 | 0.7811 | 0.7273 |
| MusNG-d15496 | 0.8830 | 0.8936 | 0.8492 | 0.9032 | 0.9681 | 0.9070 | 0.9583 | 0.9333 | 0.8075 | 0.8296 |
| MusNG-d15697 | 0.8902 | 0.8810 | 0.8727 | 0.9500 | 0.9526 | 0.9028 | 0.9810 | 0.9226 | 0.7869 | 0.9268 |
| HumN-d9861 | 0.8171 | 0.9565 | 0.5720 | 0.2830 | 009637 | 0.9167 | 0.8835 | 0.9693 | 0.6341 | 0.1744 |
| HumN-d10021 | 0..8992 | 0.9451 | 0.8191 | 0.8795 | 0.8575 | 0.9378 | 1.0000 | 0.9249 | 0.8385 | 0.9359 |
| HumN-d12876 | 0.8072 | 0.8765 | 0.5513 | 0.8810 | 0.9333 | 0.9579 | 0.8442 | 0.9181 | 0.6466 | 0.5736 |
| HumN-d14380 | 0.7675 | 0.9169 | 0.3750 | 0.8958 | 0.9844 | 0.9662 | 0.7775 | 0.9522 | 0.5118 | 0.4343 |
| HumN-d15496 | 0.7617 | 0.9468 | 0.5238 | 0.4032 | 0.9628 | 0.8750 | 0.8480 | 0.9604 | 0.5617 | 0.3356 |
| HumN-d15697 | 0.8663 | 0.9357 | 0.7182 | 0.8000 | 0.8947 | 0.9156 | 0.9667 | 0.9357 | 0.7117 | 0.8101 |
| HumNG-d9861 | 0.7600 | 0.8725 | 0.4640 | 0.7925 | 0.9967 | 0.9066 | 0.8197 | 0.9311 | 0.5383 | 0.3281 |
| HumNG-d10021 | 0.7794 | 0.7727 | 0.7553 | 0.9036 | 0.9973 | 0.8985 | 0.8346 | 0.8709 | 0.7648 | 0.5137 |
| HumNG-d12876 | 0.6915 | 0.6091 | 0.8077 | 0.9524 | 0.9583 | 0.8807 | 0.7726 | 0.7475 | 0.7200 | 0.5161 |
| HumNG-d14380 | 0.6786 | 0.5668 | 0.8611 | 0.9167 | 0.9062 | 0.9117 | 0.7547 | 0.6996 | 0.8212 | 0.4190 |
| HumNG-d15496 | 0.9234 | 0.9433 | 0.8651 | 0.9516 | 1.0000 | 0.9506 | 0.9534 | 0.9708 | 0.8651 | 0.8429 |
| HumNG-d15697 | 0.8283 | 0.7846 | 0.8727 | 0.9375 | 0.9368 | 0.9182 | 0.9002 | 0.8607 | 0.8067 | 0.7614 |

Supplementary Table S2 Each row is a combination of scRNA-Seq dataset and brain donor each corresponds to a spatial distribution figure (e.g. Figure 5B, Supplementary Figures 14-30B). The overall accuracy for k-means clusters as well as the cluster-wise sensitivity, cluster-wise specificity, and cluster-wise F-score are contained in each column.

|  | Inter-neuron | S1 Pyramidal | CA1 Pyramidal | Oligo-dendrocyte | Microglia | Endothelial | Astrocyte | Ependymal | Mural |
| --- | --- | --- | --- | --- | --- | --- | --- | --- | --- |
| MusNG-PC1 | 0.0000 | 0.8503 | 0.0043 | 0.0000 | 0.0220 | 0.0000 | 0.1234 | 0.0000 | 0.0000 |
| MusNG-PC2 | 0.1661 | 0.0000 | 0.0000 | 0.0155 | 0.0089 | 0.2815 | 0.5083 | 0.0196 | 0.0000 |
| MusNG-PC3 | 0.0484 | 0.1032 | 0.6581 | 0.0070 | 0.0062 | 0.1666 | 0.0000 | 0.0104 | 0.0000 |
| HumN-PC1 | 0.0000 | 0.6279 | 0.2020 | 0.0041 | 0.0064 | 0.0000 | 0.1596 | 0.0000 | 0.0000 |
| HumN-PC2 | 0.0000 | 0.3645 | 0.0000 | 0.0075 | 0.0000 | 0.3499 | 0.2780 | 0.0000 | 0.0000 |
| HumN-PC3 | 0.4871 | 0.0683 | 0.0180 | 0.0609 | 0.0000 | 0.0145 | 0.3507 | 0.0005 | 0.0000 |
| HumNG-PC1 | 0.6156* | 0.6156* | 0.0680 | 0.0000 | 0.1918 | 0.0000 | 0.0000 | 0.1245 | 0.0000 |
| HumNG-PC2 | 0.0075* | 0.0075* | 0.0747 | 0.0301 | 0.0584 | 0.6839 | 0.0314 | 0.1136 | 0.0003 |
| HumNG-PC3 | 0.0899* | 0.0899* | 0.0000 | 0.4358 | 0.0000 | 0.0180 | 0.0060 | 0.4503 | 0.0000 |

Supplementary Table S3 Cell type makeup of each of the 3 principal cell types used to generate Figure 7. * In the HumNG dataset there are only 8 cell types because the interneuron and S1 pyramidal were combined. The percentages in those cells are for the combined cell type.
